# Characterization of the p38α MAPK allosteric inhibition by a single chain Fv antibody

**DOI:** 10.64898/2026.01.06.697868

**Authors:** Emilie Renaud, Marion Larroque, Nicole Bec, Hugo Guérin, Raphaël Sierocki, Corinne Henriquet, Martine Pugnière, Oriane Font, Alexandra Fauvre, Vincent Denis, Céline Gongora, Christian Larroque, Pierre Martineau, Laurence Guglielmi

## Abstract

Most of the available p38α kinase inhibitors target the highly conserved ATP-binding site, thus explaining their off-target effects and toxicity. Here, we report the use of single-chain fragment variable (scFv) antibodies to identify new highly specific inhibitors. Using phage display, we selected five anti-p38α scFv antibodies among which one fully inhibited p38α kinase activity *in vitro*. We showed that this scFv did not affect ATP binding, while still acting as a competitive inhibitor of ATP hydrolysis. When expressed as intrabody in the nucleus of the THP-1 human monocytic cell line, the scFv inhibited the function of endogenous p38α and induced a significant decrease in lipopolysaccharide-induced tumor necrosis factor α production. To gain insight into the mechanisms by which the scFv induced p38α inhibition, the scFv epitope was mapped by deep mutational scanning of p38α displayed on yeast surface and high-throughput selection by flow cytometry of p38α mutants no longer bound by the scFv. By this approach, we showed that the inhibitory scFv interacts with two helices within the C-terminal lobe of the kinase distant from the ATP-binding pocket and the docking groove. Altogether, our data show that the scFv antibody behaves as an allosteric inhibitor, constraining and maintaining p38α in an inactive form.

## Introduction

Overexpression or dysregulation of protein kinases have been directly associated with cancer and many human diseases due to their involvement in important and essential physiological responses of eukaryotic cells such as proliferation, apoptosis and cell signaling. As a result, protein kinases are one of the most popular classes of drug targets. The human kinome comprises more than 500 kinases, most of which share a common structure made of two lobes (N-terminal and C-terminal), separated by a cleft that contains the highly conserved ATP binding pocket comprising residues important for ATP-binding as well as phosphate transfer. Kinase inhibitors are classified based on their binding site and the structure of the drug-enzyme complex^1^. Type I and II inhibitors are ATP competitors that bind within the ATP pocket. While type I inhibitors alter the active state of the enzyme, type II inhibitors bind to the inactive conformation of the kinase. Type III and IV are allosteric inhibitors that do not target the ATP pocket. Type III inhibitors bind to a pocket adjacent to the ATP site, whereas type IV inhibitors bind to a site distant from the ATP pocket without blocking ATP or substrate binding sites. Type V are bivalent inhibitors that bind to two different sites of the kinase. Lastly, unlike the type I-V inhibitors that are reversible, type VI are suicide inhibitors that form an irreversible covalent bond with the targeted kinase.

Drug specificity has rapidly emerged as an important issue. Indeed, the majority of the available kinase inhibitors act as ATP competitors, explaining why most of them are associated with unrelated off-target kinase inhibition^2^. These off-target effects partly explain the lack of efficiency and the toxicity and are an obstacle to the development of new drugs. Therefore, efforts have now shifted to the identification of allosteric inhibitors aimed at exploiting structural features and regulatory mechanisms that are unique to a particular kinase. However, among the 74 currently FDA-approved small molecule protein kinase inhibitors, only three are classified as type III and five as type IV^3^.

In the present study, we report a new strategy to target the p38α kinase by the use of antibody fragment. The p38 mitogen-activated protein kinase (MAPK) family include four members (α, β, γ, δ) that share more than 60% amino acid sequence identity. These serine-threonine kinases respond to a wide range of extracellular stimuli, essentially environmental stress like UV radiation, osmotic shock, oxidative stress, inflammatory cytokines and hypoxia^4,5^. In the p38 MAPK family, p38α (or MAPK14) is the best-studied isoform considering that its ubiquitous and strong expression. Due to its role in inflammatory, neurodegenerative and autoimmune diseases as well as cancer, p38α has become a therapeutic target of interest and significant efforts have been made to develop selective inhibitors of its kinase activity. Despite all these efforts, none of these molecules have been approved for clinical use.

The structure of p38α is well known, thanks to the hundreds of p38 crystal structures available (250 PDB entries). Conformational changes are required for full p38α kinase activity^6^. Stress-activated MAPK Kinases (MKK) phosphorylate the threonine and tyrosine residues in the TGY motif located in p38α activation loop. This dual phosphorylation induces a conformational change of the activation loop that brings the N- and C-terminal lobes closer, facilitating the reorientation of key residues for ATP and Mg2^+^ ion stabilization. In addition, p38α C-lobe contains a docking groove composed by the glutamate-aspartate (ED) domain^7^ and a common docking (CD) motif for interactions with the kinase interaction motif (KIM or D-motif) of upstream activators, downstream substrates, and phosphatase regulators^8^. The study of the dynamics of p38α activation by NMR demonstrated that both phosphorylation and KIM binding are necessary to change p38α into an activated state^9^.

Here, we present an approach to target p38α using single chain Fragment variable (scFv) that are small antibody fragments (28 kDa) carrying a full antigen binding site composed of the variable domains of the heavy (VH) and the light chain (VL) joined by a flexible linker. ScFvs combine high specificity and affinity for their antigen and are able to distinguish closely related family members and to target a specific protein conformation or post-translational modification^10–14^. They are powerful tools for targeting cytoplasmic proteins through their ability to directly inhibit protein activity or protein-protein interactions^10,11,15,16^. In this study, we successfully selected a scFv specific for the p38α isoform that can fully inhibit the kinase activity *in vitro*. We mapped the scFv binding site on p38α and showed that it does not block the ATP and substrate binding sites and inhibits p38α kinase activity probably by maintaining the enzyme in an inactive conformation.

## Results and Discussion

### Identification of anti-p38α antibody fragments

To isolate p38α highly specific scFvs, we performed a phage display selection after depleting our antibody fragment library^17,18^ of the cross-reacting scFvs able to bind to p38β, which shares 74% sequence identity with p38α. After three rounds of biopanning, five different scFvs, namely B2, B12, D9, E5 and G9, were identified. We checked their specificity and showed that all scFvs selectively bound p38α by ELISA (Figures S1, A and B) and immunoprecipitated endogenous p38α from HCT116 cell protein extracts (Figure S1C). Sequence analysis showed that scFvs B2 and E5 present similar complementary determining regions (CDR), as well as B12 and D9, while G9 differ from the other scFvs in its amino acid composition (Figure S2A). These data suggested that at least three dependent p38α epitopes were targeted. To confirm this hypothesis, the scFvs were tested in competitive ELISA assays for p38α binding (Figure S2B). The results showed that G9 bound to a distinct epitope from the other scFvs, while B2 and E5 on one hand and B12 and D9 on the other hand shared an overlapping binding site.

### The scFv G9 is a specific inhibitor of p38α kinase activity

There are different ways to block p38α. One is to inhibit p38α activation, which is achieved by the dual phosphorylation of its Thr_180_ and Tyr_182_ residues by the MAPK kinases MKK3 and MKK6. Another is to inhibit directly p38α kinase activity. To identify a p38α inhibitor among our five scFvs, we first performed an *in vitro* kinase assay to test their ability to block p38α phosphorylation by a constitutively active MKK6. The results showed that all scFvs significantly decreased p38α phosphorylation, with an inhibition ranging from 65% to 85% depending on the scFv (Figure 1A). Since the different scFvs bound non-overlapping regions of p38α (Figure S2B), we hypothesized that they probably act individually through a steric inhibition mechanism by preventing MKK6-p38α interaction.

**Figure 1.**
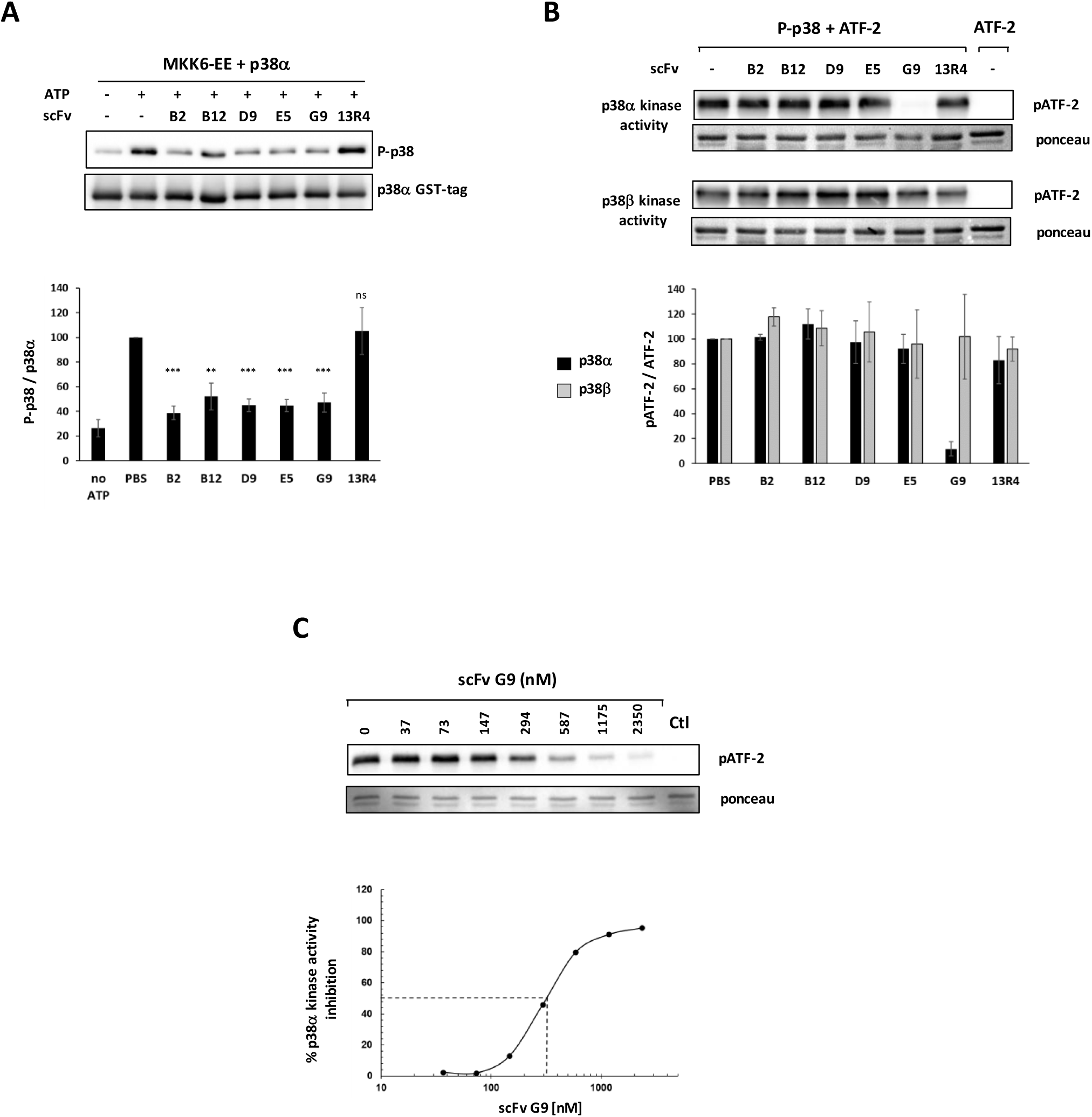
Inhibition of p38α kinase activity by scFv. **(A)** *In vitro* p38α phosphorylation by MKK6. Kinase reactions containing p38α, a constitutively active mutant of MKK6 (MKK6-EE), and the indicated anti-p38α scFv were incubated at 22°C for 14h and analyzed by western blotting. An anti-GST tag antibody was used to detect total p38α. The anti-β-galactosidase 13R4 scFv was used as irrelevant antibody. Lower panel: band quantification (mean ± SD of 4 independent experiments). **p < 0.01, ***p < 0.001 (Student’s *t*-test). **(B)** Western blot analysis of ATF-2_(19-96)_ phosphorylation by activated P-p38α or P-p38β in the presence of the indicated anti-p38α scFv. Reactions were stopped after 30-min incubation at 30°C. Lower panel: band quantification (mean ± SEM of 3 independent experiments). For A and B: p38α/scFv molar ratio was 1:2.5. **(C)** Dose-response curve of p38α inhibition by G9, determined by Western blot analysis and ATF-2_(19-96)_ phosphorylation quantification. An irrelevant scFv was added to each point of the reaction to maintain an equal amount of scFv. For B and C, the last lane corresponds to a control without kinase.

We then monitored the effect of each scFv on p38α activity using another kinase assay that follows p38α or p38β activity on their common target ATF2. For this purpose, strongly phosphorylated and activated p38 proteins (P-p38) were produced^19^. The results showed that G9 was the only scFv able to abolish almost completely ATF2 phosphorylation by p38α and that G9 inhibits p38α kinase activity in a dose-dependent manner Figure 1, B and C). As expected, G9 had no effect on p38β activity. None of the other four scFvs were able to alter ATF2 phosphorylation by p38α or by p38β.

### The scFv G9 does not bind the ATP pocket, but inhibits ATP hydrolysis

To gain insights into the mechanisms by which G9 inhibits p38α enzymatic activity, we sought to compare G9 with SB 202190, a potent type I inhibitor of both p38α and p38β isoforms. The inhibitory potential of these compounds was evaluated using the ADP-Glo kinase assay that measures ATP hydrolysis. We used as substrate a short 16-amino acid peptide that contains serine-threonine protein kinase phosphorylation sites. By this approach, we found that G9 inhibited p38α at nanomolar concentrations and IC_50_ values for G9 and SB 202190 were estimated respectively at 150 nM and 30 nM (Figure 2A). We also confirmed the specific inhibitory effect of G9 on p38α kinase activity, as G9 at high concentration could inhibit only 25% of p38β kinase activity (Figure S3). We then tested G9 and SB 202190 in combination (Figure 2B, Figure S4). In the presence of SB 202190, we observed a two-fold decrease of G9 IC_50_ (75 nM). The inhibition curve fitted perfectly with the curve predicted by the Bliss’ model that stipulates the independence of the two inhibitors^20^. These results suggested that G9 and SB 202190 act by different and independent mechanisms on p38α activity and exert additive inhibitory effects.

**Figure 2.**
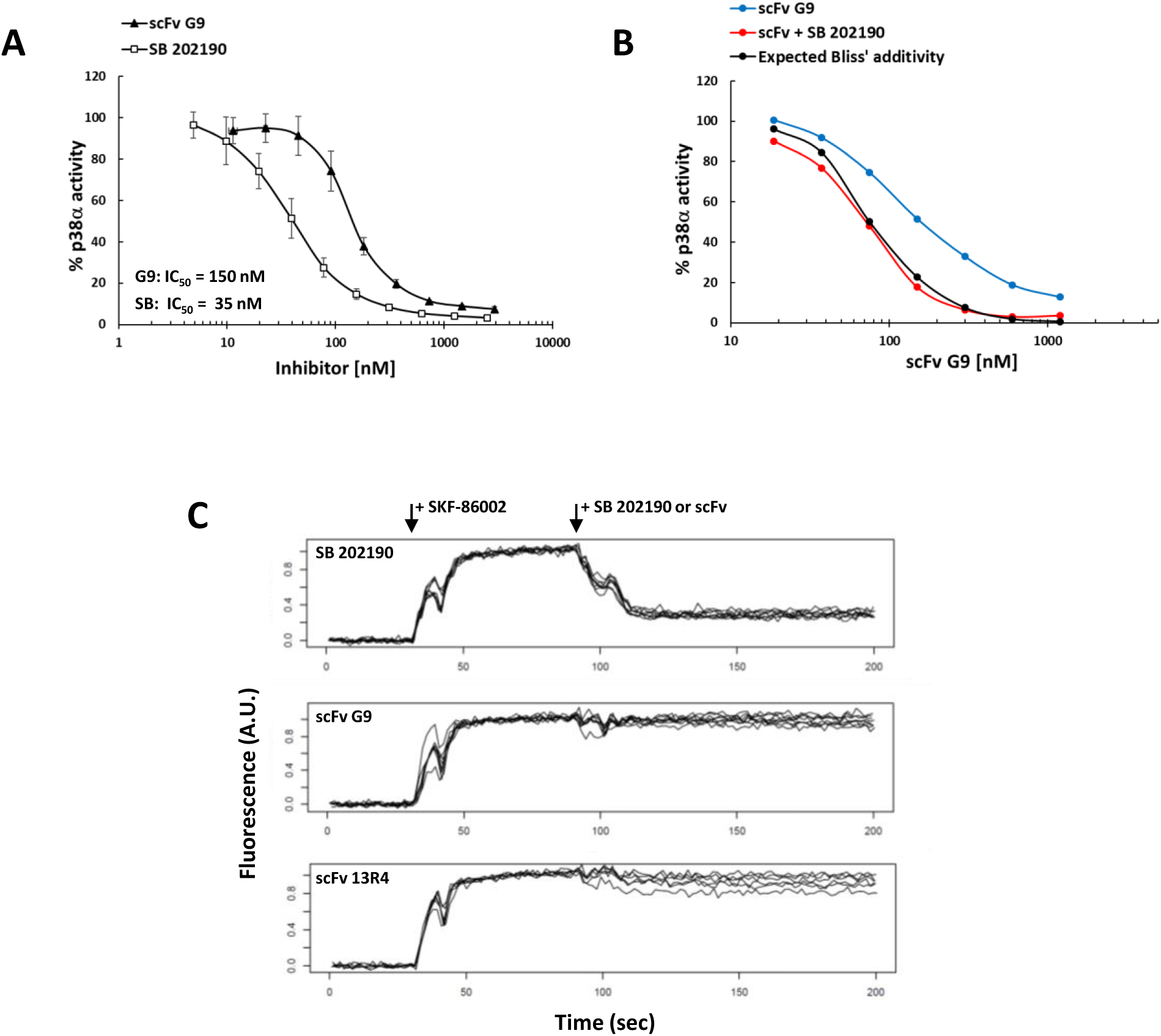
Comparison of inhibitory and binding activities of the scFv G9 and the pyridinyl imidazole p38 inhibitor SB 202190. **(A-B)** The *in vitro* kinase inhibition potency of p38α inhibitors was measured in ADP-Glo assays using as substrate a synthetic peptide (IPTTPITTTYFFFKKK) that contains Ser and Thr phosphorylation sites of kinase proteins. After preincubation of the inhibitors with p38α for 20 min, reactions were started by adding ATP, and stopped after 1h incubation at room temperature. **(A)** Dose-response inhibition of p38α activity by the scFv G9 and SB 202190 (n=3 independent experiments). **(B)** Combination of scFv G9 and SB 202190. The black line represents the percentage of p38α kinase activity calculated according to the Bliss’ model assuming the independence of the two inhibitors. **(C)** Association curves for the binding of SKF-86002 to p38α, determined by real-time measurements of the ATP-competitive inhibitor SKF-86002 fluorescence emission at 420 nm. The excitation wavelength was 340 nm. SKF-86002 was first added to a p38α solution at time 30 sec and then SB 202190, scFv G9 or scFv 13R4 (irrelevant) were added at time 90 sec. The decrease in fluorescence signal corresponds to SKF86002 displacement from the p38α ATP binding site. The concentrations used are: p38α 200 nM; SKF86002 200 nM; SB202190 2 µM; scFv 1 µM. For each compound, n=8. Time-resolved fluorescence was measured using the FLEXstation 3 reader.

SKF86002 is a p38α inhibitor that lacks intrinsic fluorescence, but generates fluorescence upon binding to the ATP-binding sites of p38α with an estimated Kd of 180 nM^21^. SKF86002 was used previously to screen ATP-competitive inhibitors, allowing the selection of new compounds that bind to p38α with a better Kd and that can displace SKF86002 from the ATP pocket^22^. Surface plasmon resonance experiments showed that G9 had a good affinity, similar for p38α and P-p38α (Kd 75 ± 6.9 nM and 78.9 ± 2.9 nM, respectively), that could enable G9 to displace SKF86002 from the p38α ATP pocket (Figure S5). Thus, we performed a displacement binding assay using SKF86002 (Figure 2C) and showed that unlike SB 202190, G9 binding to p38α did not lead to a decrease in the fluorescence signal (<1% vs. 70% for SB 202190). Furthermore, SKF86002 was still able to bind p38α when complexed to G9 (Figure S6C). This indicated that G9 does not act as an ATP competitive ligand, does not block ATP and analogs binding and probably binds to a site distinct from that of inhibitors targeting the kinase domain.

We then studied the inhibitory activity of G9 at various ATP concentrations using the ADP-Glo assay. The resulting Lineweaver-Burk plot showed that G9 behaved as a competitive inhibitor of ATP hydrolysis, although it did not block ATP binding (Figure 3B). Indeed, G9 induced an increase of p38α Km without modifying its Vmax. The inhibition constant (Ki ∼37 nM) was in the same order of magnitude as that of chemical inhibitors of p38 (Figure 3C). Using the same experimental approach, we showed in contrast that G9 was not a competitive inhibitor of the substrate peptide (Figure 3A). Therefore, we hypothesized that G9 could inhibit p38α activity not by blocking its interaction with its substrates, but by inducing a conformational change that stabilizes the enzyme in an inactive conformation.

**Figure 3.**
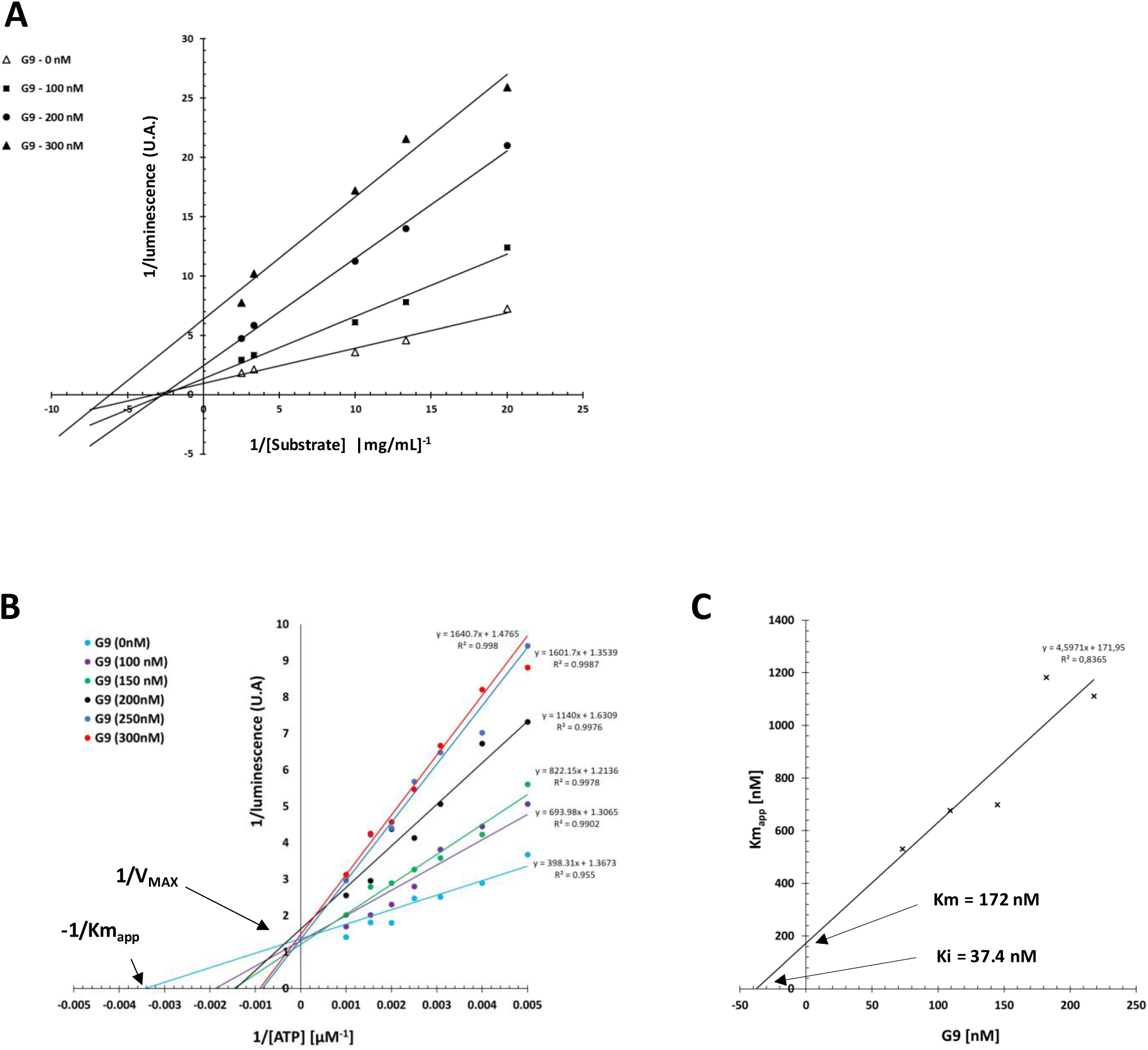
scFV G9 is a competitive inhibitor of ATP hydrolysis. **(A-B)** Lineweaver-Burk plots: curves represent ATP hydrolysis by p38α. The ATP-Glo assay was performed using 26 µM p38α and different concentrations of G9 (0, 100, 150, 200, 250 and 300 nM) for each substrate peptide IPTTPITTTYFFFKKK concentration (0.4, 0.3, 0.1, 0.075 and 0.05 mg/mL) **(A)** or for each ATP concentration (50, 100, 200, 250, 325, 400, 500, 650, 1000 µM) **(B).** After preincubation of the scFv G9 with p38α for 20 min, the reaction was started by adding ATP and stopped after 30-min incubation at room temperature. **(C)** Determination of scFv G9 inhibition constant (Ki) against ATP. The apparent Km (Km_app_) was calculated for each G9 concentration on the Lineweaver-Burk plot B.

### Identification of G9-p38α interaction interface

To better understand G9 mechanism of inhibition, we determined the epitope recognized by the scFv. Currently, the gold standard for epitope mapping is to solve the atomic structure of antigen-antibody dimer crystals by X-ray diffraction. This approach would have allowed measuring potential p38α conformational changes when bound to G9. Unfortunately, the scFv G9 aggregates at high concentrations, thus hindering crystal formation. Therefore, we used a high throughput approach consisting in a combination of yeast surface display (YSD) and mutational scanning (DMS) to identify which p38α amino-acids are involved in scFv G9 binding^23,24^. Briefly, a point mutant library was generated in which each of the 360 amino acids of p38α was substituted by all of the other 19 natural amino acids. Expressed on the surface of yeast cells, mutants with altered antibody recognition were enriched by fluorescence-activated cell sorting (FACS). Next generation sequencing (NGS) of plasmids extracted from sorted population then allowed to identify binding disruptive mutations. Furthermore, this approach allowed simultaneous mapping of G9 and B2 epitopes. Indeed, since these scFvs do not compete for p38α binding, yeasts could be co-stained with both scFvs. To avoid selecting mutants that could not be displayed on yeast or were misfolded, our strategy was to isolate by FACS cells that lost binding for only one antibody but not the other (Figure 4A).

**Figure 4.**
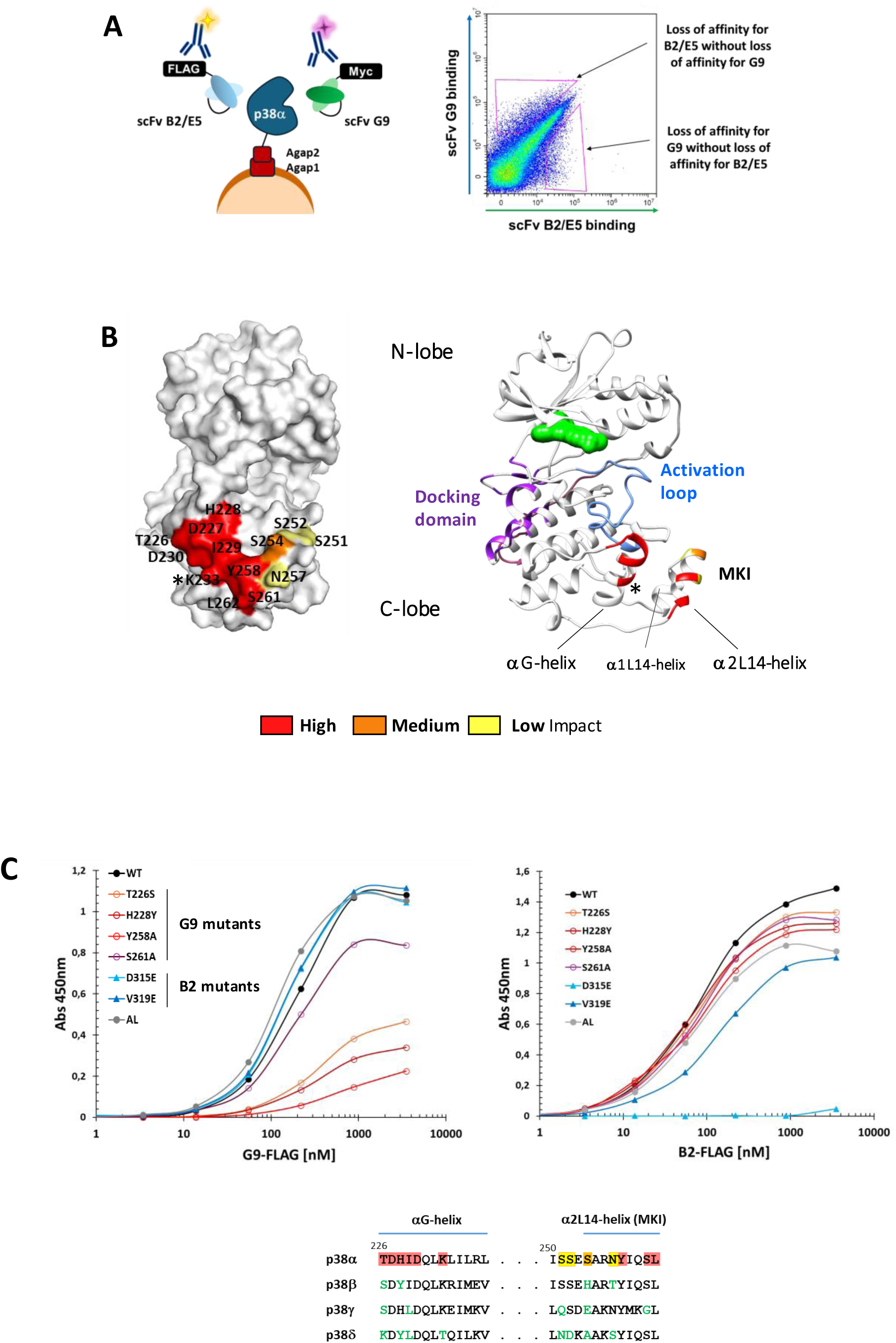
p38α Epitope mapping. (A) Schematic of functional screening of scFv G9 and scFv B2/E5 binding to p38α mutants by yeast surface display. The right panel shows the gating strategy for FACS enrichment of mutations that impaired scFv binding. (B) Structure of p38α as a surface (left) and a ribbon (right) showing scFv G9 epitope. The residues involved are colored from red (high) to yellow (low) to highlight their impact on the interaction. The docking domain (purple) and the activation loop (blue) are indicated. As a reference, an ATP competitor is shown as a green surface in the ATP binding pocket. The asterisk (*) indicates K233 residue involved in DSBU cross-linking of scFv G9-p38α complex. (C) Epitope validating by ELISA binding assay. ELISA were performed by testing the binding of FLAG-tagged scFvs to immobilized wild-type or mutant p38α. The anti-FLAG antibody conjugated to HRP was used for detection.

Sequence data were represented as heatmaps (Supplementary Table S1). Analysis of mutants escaping G9 binding led us to identify key epitope residues, all located at the surface of two helices of p38α C-lobe: T226, D227, H228, I229, D230, K233 in the αG-helix and Y258, S261, L262 in the α2L14-helix of the MAPK insert (MKI) (Figure 4B). Two other amino acids located in the α2L14-helix had medium (S254) or low (N257) impact on G9 binding when mutated. Residues S251 and S252 located in the loop connecting the two MKI helices (α1L14 and α2L14) had only low impact. All together, these results show that scFv G9 recognizes a conformational epitope.

On the other hand, scFv B2 also binds to a conformational epitope in the C-lobe of p38α, but on the opposite side of the protein. The epitope comprises ten amino acids whose substitution strongly impaired B2 binding (Figure S7). Eight amino acids (A304, A309, Q310, H312, D313, P314, D315, D316, V319) are located in the loop containing the docking domain for substrates or regulators of p38α activity. The two other amino acids (D125 and F129) are present in the αE-helix. Finally, the functional heatmap indicated that substitution of K79 residue had medium impact on B2 binding. As a control, we also performed scFv E5 epitope mapping and confirmed the results of competitive binding assays showing that the B2 and E5 scFvs shared overlapping epitopes. Indeed, all key residues for B2 binding were also implicated in E5 binding.

### Epitope validation

To validate results of DMS experiments, we produced a selection of recombinant mutants and tested scFv binding by ELISA. Point mutations were chosen according to the heatmap (Supplementary Table S1) and corresponded to substitutions with the amino acid of the p38β sequence when different or to an alanine when identical. For G9 epitope validation, we chose two mutations within the αG-helix (T226S and H228Y) and two mutations within the α2L14-helix (Y258A and S261A). As controls, we produced two mutants of the B2 epitope (D315E and V319E) and a mutant (AL) carrying the p38β activation loop sequence (H174Q-T175A-D177E).

Compared to the wild type p38α, each of the T226S, H228Y and Y258A substitutions greatly reduced binding for G9, but not for B2, confirming that these residues play an important role in G9 recognition without globally affecting p38α structure (Figure 4C). In contrast, we could not confirm the key role of S261 as its substitution caused only modest reduction in G9 binding. As expected, the other substitutions had no impact. Overall, these results were consistent with the G9 epitope identified by the yeast-display DMS approach. We also confirmed the region of interaction of scFv B2 with p38α. Indeed, despite a low impact of the V319E substitution on B2 binding activity, the D315E mutation induced a total loss of recognition of B2 while retaining its affinity for G9.

To further validate the epitope, we performed a Mass Spectrometry (MS) analysis of the G9-p38α complex, chemically bound with the MS-cleavable cross-linker disuccinimidyl dibutyric urea (DSBU). Cross-linking mass spectrometry (XL-MS) has become a powerful approach to provide low resolution structural information on proteins and protein-protein interactions^25,26^. We chose the DSBU reagent, a medium long-spaced arm suitable for intermolecular cross-linking that creates covalent bounds between lysine residues with a distance constraint of 26-30 Å between Cα atoms^27,28^.

First, we tested different DSBU concentrations to optimize cross-linking reactions. Proteins were diluted at 10 µM at a molar ratio of 1:1 and incubated 1h at 25°C with various molar excess of DSBU. Reactions were analyzed by SDS-PAGE (Figure S8A). At 5-fold molar concentration of DSBU (50 μM), we observed two main protein complexes at approximately 70 and 95 kDa. Both contained G9 (27 kDa) and p38α (42 kDa), as indicated by western blotting using anti-MYC-tag and anti-p38α antibodies (Figure S8B, lanes 2 and 10). We concluded that the p38α-G9 complex existed in two stoichiometries: 1 p38α:1 G9 and 1 p38α:2 G9. This result was not surprising as it is well documented that scFvs can form active dimers (i.e. diabodies)^29^. This was further confirmed by the additional band at 60kDa that corresponded to the G9 diabody (lane 2), also observed in the absence of p38α (lane 4). The 95 kDa band presumably contained some p38α dimers because it was present also in the absence of G9, although at much lower intensity (lanes 11 and 14). Lastly, in the absence of DSBU or when G9 or p38α was replaced by bovine serum albumin (BSA), we did not detect any complex (lanes 1 and 9, 6 and 14, and 8 and 16, respectively), demonstrating the specificity of the cross-link. The same SDS-PAGE migration profile was observed for the B2-p38α cross-linked complex (date not shown).

For XL-MS, we used 100-fold molar excess of DSBU (1 mM) as most of the proteins were cross-linked at this concentration, and analyzed the band at 95 kDa, which corresponds to the complex of interest, the diffuse pattern being probably due to intramolecular cross-links that modify protein migration. The MeroX software was used to scan MS/MS spectra^30^ and one major intermolecular cross-links was identified for each scFv-p38α complex: between K64 of G9 VH domain (according to Kabat numbering scheme) and K233 of p38α, and between K64 of B2 VH domain and K79 of p38α (cross-linked Lys are indicated by an asterisk in Figures 4B, 4C and S7A). These results are consistent with our previous observations, as K233 and K79 are important residues found by the yeast display DMS experiment as part of G9 and B2 epitopes, respectively.

### scFv G9 binds to an allosteric site

Epitope mapping represents a great challenge. The combination of DMS, yeast display and high throughput sequencing allowed to determine unambiguously the interaction interface between G9 and p38α. The scFv G9 interacts with p38α through its C-lobe without blocking the ATP binding site or the common docking site (Figure 4B). These results are consistent with our experimental data, demonstrating that G9 is not an ATP competitive inhibitor, and suggest that G9 blocks ATP hydrolysis by restraining the flexibility of p38α, a key characteristic of the enzyme. Our hypothesis is supported by several studies showing that the region targeted by our antibody fragment contains an allosteric site allowing modulation of p38α activity. Indeed, the αG helix and both helices of the MAPK insert (α2L14 and α1L14 helix) form a hydrophobic pocket, also called lipid-binding domain (LBP)^31^, that has been shown to be important for p38α specific interaction with the substrate ATF2^32^ or the activator MKK6^33^. NMR and crystallographic studies highlighted that ligands bound to this cleft directly modulate p38α activity by inducing an allosteric structural change^31,34–36^. To our knowledge, only one allosteric inhibitor targeting p38α pocket has been identified. Selected by screening small molecule libraries, Compound 10 described by Comess et al.^34^, exhibits high selectivity for the p38α isoform within the p38 MAPK family. However, analysis of the binding sites of scFv G9 and compound 10 revealed that they are distant and clearly distinct from each other. Our findings therefore indicate that the LBP pocket is not the only region within the C-lobe of p38α that can be targeted, but that neighboring sites also enable modulation of p38α kinase activity. Altogether, our results show that the scFv G9 acts as an allosteric inhibitor of p38α and that the use of antibody fragments provides an interesting approach to find new pharmacological targeting strategies. Unlike small chemical molecules that need to fit into pockets to bind stably to proteins, antibodies interact with proteins across flat surfaces. Thus, the selection and characterization of a p38α-specific scFv could allow the identification of a new class of allosteric inhibitors, different from those already characterized.

### The G9 intrabody binds to p38α in cells

ScFvs can be easily expressed in mammalian cells as functional intracellular antibodies, also called intrabodies. To verify whether G9 can bind specifically to its target also in cells, we performed a retargeting assay^37^. ScFv-mCherry fusion protein were co-expressed in transiently transfected Hela cells with GFP-p38α or GFP-p38β fusion proteins harboring a nuclear localization signal (NLS) or a nuclear export signal (NES). Fluorescence microscopy analysis showed that G9 presented a high soluble expression level and conserved its specific binding activity within the cell. Indeed, when G9 and p38α were over-expressed together, the antigen and the antibody were detected in the cytoplasm and the nucleus, whereas p38α expression with an NLS or an NES induced a re-localization of G9 in the nucleus or the cytoplasm, respectively (Figure 5). Conversely, p38β re-localization did not induce a change of cellular distribution of G9, indicating that the intrabody did not bind to p38β in HeLa cells.

**Figure 5.**
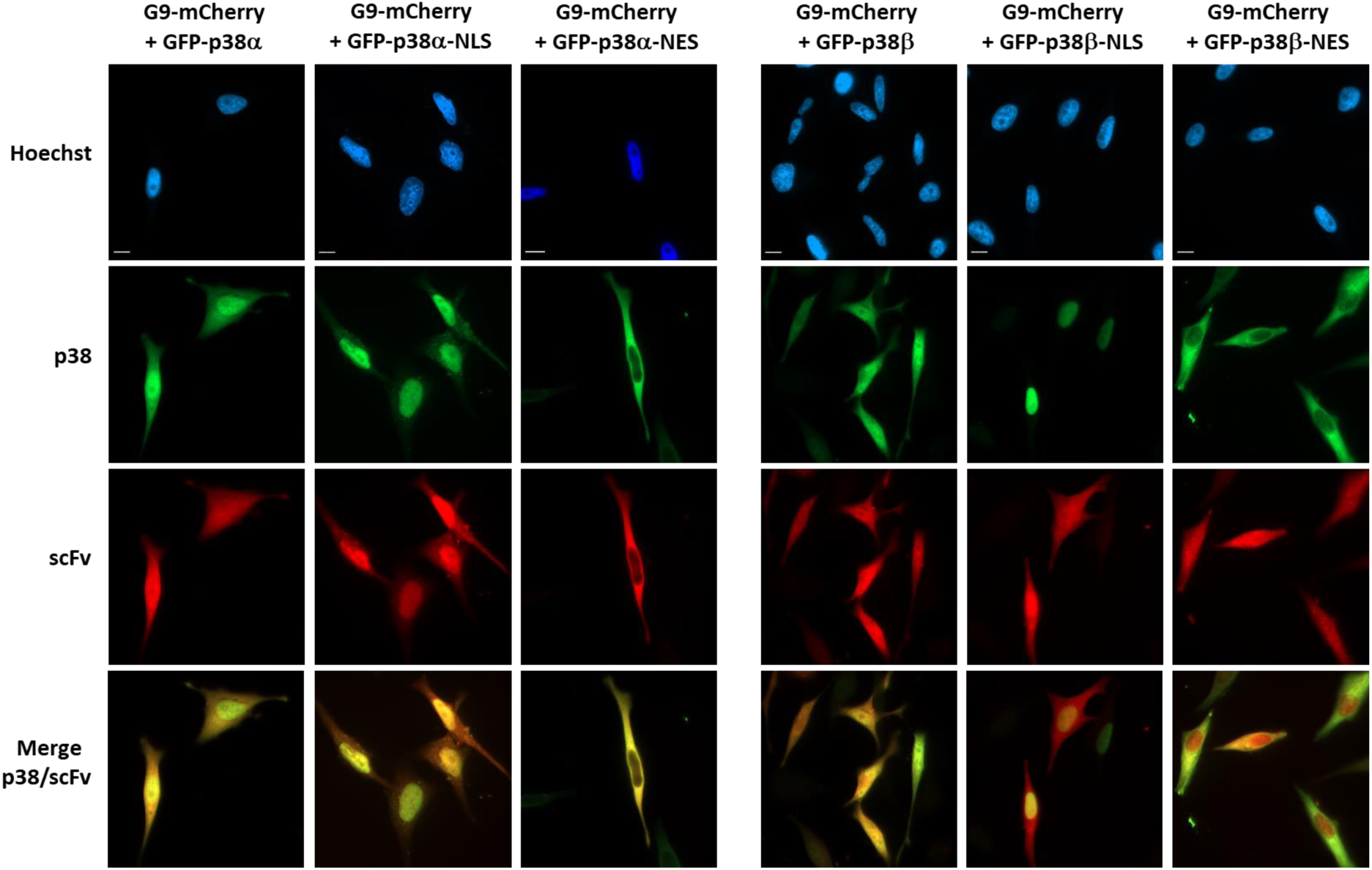
Interaction of G9 intrabody with p38α in HeLa cells. Analysis of the subcellular distribution of G9-mCherry intrabody in HeLa cells co-expressing NLS- or NES-tagged GFP-p38α or GFP-p38β. Cells were fixed 48h after transient transfection and DNA was stained with Hoechst. Fluorescence was examined with a ZEISS Axio Imager Z2 microscope (X63 objective). Scale bars represent 10 μm.

As a control for the next experiments, we needed a scFv that could bind to p38α, but without blocking its enzymatic activity. Therefore, we also tested the cell expression and binding properties of the other four anti-p38α scFvs selected by phage display. Only B2 was strongly expressed and specifically bound to p38α (Figure S9A). Conversely, B12, D9 and E5 were poorly expressed or formed aggregates, indicating their inability to fold correctly in the intracellular reducing environment (data not shown). In line with these results, their capacity to bind to p38α in cells was reduced.

Altogether, these results suggest that the G9 and B2 intrabodies had sufficient expression and affinity to bind to p38α within cells without evidence of protein aggregation.

### The G9 intrabody inhibits LPS-induced TNF-α production by THP-1 cells

Several p38 inhibitors have been tested for their ability to inhibit lipopolysaccharide (LPS)-induced TNF-α production by monocytes. Indeed, upon LPS stimulation, p38α is activated and translocates into the nucleus where it phosphorylates MAPK-activated kinase 2, an essential protein that regulates TNF-α biosynthesis^38–40^. We therefore used this phenotypic assay to investigate whether G9 can modulate the endogenous p38α function in cells. The human monocytic cell line THP-1 was transduced for an inducible expression of the G9 or B2 intrabodies fused to GFP and a NLS tag to target p38α nuclear function. Before LPS stimulation, cells were differentiated into macrophages by incubation with phorbol 12-mystyrate 13-acetate (PMA) for 48h and concomitantly treated with doxycycline that induced intrabody expression in an average of 79% of cells, as monitored by flow cytometry (Figure 6A). Immunoblot analysis performed using nuclear and cytoplasmic protein extracts of differentiated THP-1 cells stimulated with LPS for 30 min indicated that G9-GFP-NLS intrabody was mainly present in the nucleus where the phosphorylated p38α pool also was found (Figure 6B). Conversely, B2-GFP-NLS was mostly maintained in the cytoplasm and the NLS failed to ensure localization of the GFP alone in the nucleus.

**Figure 6.**
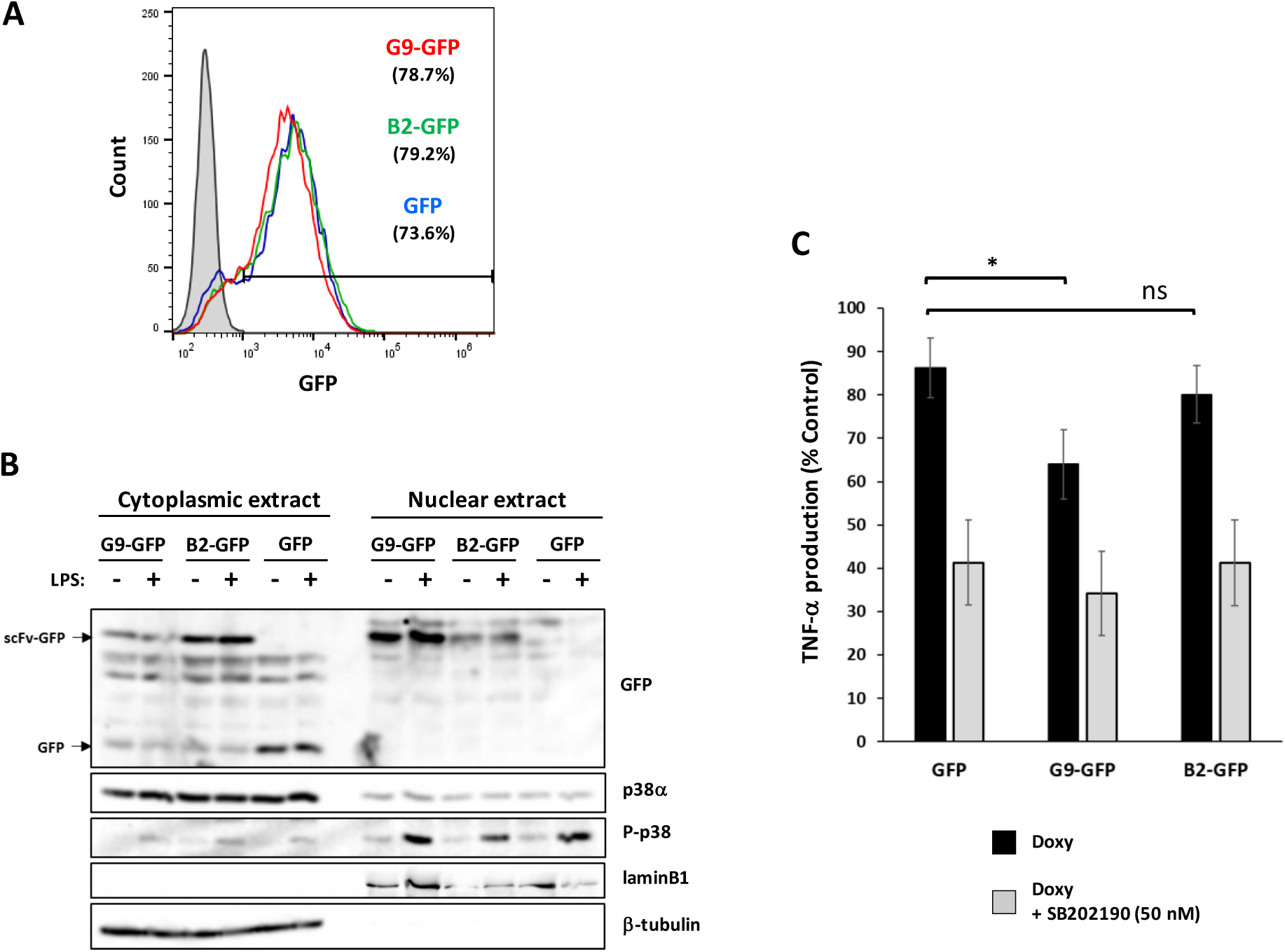
LPS induction of TNF-α expression in THP-1 cells is inhibited by G9 intrabody. Transduced THP-1 cells were treated for 48h with PMA (10 ng/mL) to induce differentiation into macrophages and with doxycycline (1 µg/mL) to induce scFv-GFP-NLS expression. As control, THP-1 cells were transduced with empty vector leading to the expression of the GFP-NLS protein. (A) Flow cytometry analysis of GFP-scFv expression in differentiated THP-1 cells. **(B)** Immunoblot analysis of intrabody expression (GFP) and p38α phosphorylation (P-p38α) in cytoplasmic and nuclear fractions (loading ratio: 1:6.5) prepared from cells stimulated or not with LPS (5 ng/ml) for 30 min. **(C)** TNF-α secretion in culture supernatants of differentiated THP-1 cells stimulated with LPS (5 ng/mL) for 4h was determined in duplicate by ELISA. As control, THP-1 were also treated with SB 2029190 (50 nM) during LPS stimulation. Data were normalized to doxycycline-untreated control (THP-1) and shown as mean ± SD of 4 independent experiments. *p <0.05 (Student’s *t*-test).

To examine p38α inhibition by G9 intrabody, we stimulated differentiated THP-1 cells with LPS for 4h and then measured TNF-α production by ELISA. In THP-1 cells that stably express G9-GFP, TNF-α secretion was weakly but significantly decreased compared with cells that express GFP (22% inhibition) or B2-GFP intrabodies (6% inhibition) (Figure 6C). Treatment of cells with 50 nM SB202190 (corresponding to the IC₅₀ concentration), intended to reduce the proportion of active p38α, did not further enhance the apparent inhibitory activity of the intrabody G9. A plausible explanation for the difference observed between the *in vitro* and in cell inhibitory effects of G9 is that G9 binds with comparable affinity to both the active and inactive forms of p38α (Fig. S5), as well as to the SB202190-bound complex (data not shown). Moreover, the expression level of G9 in THP-1 cells may not be sufficient to fully block p38α activity in the nucleus, and the remaining kinase activity may be sufficient to sustain significant TNF-α secretion. Nevertheless, our results show that the G9 intrabody is able to inhibit p38α function in cells as *in vitro*.

## Conclusion

In conclusion, we identified a potent and highly selective protein inhibitor of p38α. The scFv G9 antibody fragment binds to a site located in the C-terminal lobe of p38α that is distinct from the ATP-binding pocket, demonstrating the potential of targeting allosteric regions to modulate kinase activity with high specificity. Such a mechanism may offer significant advantages over conventional ATP-competitive inhibitors, which often lack selectivity due to the high conservation of the ATP-binding site across the kinome. Beyond its allosteric inhibitory activity, the scFv may also interfere with protein-protein interactions as its epitope overlaps with a key region involved in binding substrates, activators, and regulators partners of p38α. Thus, by recognizing both active and inactive conformations of p38α, scFv G9 could inhibit not only its activation by MAPK kinases but also its catalytic function.

If the identification of scFv G9 provides new insights into the mechanisms regulating MAP kinase activity, it also opens promising avenues for the development of novel therapeutic strategies. Although intrabodies are not readily applicable as therapeutic agents, scFvs can be used to select small molecules. Indeed, our laboratory previously reported the isolation, using an antibody displacement assay, of a drug candidate that mimics the inhibitory effect of an scFv directed against the cytoplasmic tyrosine kinase SYK and that was used *in vivo* to prevent anaphylactic shock in mice^41,42^. Subsequently, this same approach was used to isolate small molecule inhibitors of a mutated form of KRAS that is frequently detected in human cancers^15^. We therefore plan to select cell-permeant peptidomimetics of the scFv G9. They can be easily tested for their capacity to inhibit p38α function in cell-based assays and will allow structural studies to better understand p38α allosteric inhibition. Lastly, the use of antibodies to identify new pharmacological targeting strategies suggests the possibility of exploiting these tools to develop inhibitors against cytoplasmic proteins classified as undruggable.

## Experimental procedures

### Reagents and antibodies

Antibodies against p38α, p38β, phospho-p38 (Thr180/Tyr182), phospho-ATF2 (Thr71), Glutathione-S-transferase (GST) and β-tubulin were purchased from Cell Signaling Technologies. Anti-FLAG M2 and anti-poly-Histidine antibodies were obtained from Sigma-Aldrich, anti-GFP and anti-c-MYC (clone 9E10) antibodies from Santa-Cruz Biotechnology.

The recombinant GST-ATF2_(19-96)_ fusion protein and the constitutively active mutant MKK6-EE were purchased from BPS Bioscience and Merck Millipore, respectively. The synthetic peptide (IPTTPITTTYFFFKKK) used as p38 substrate was from SignalChem.

PMA, LPS, doxycycline, SB 202190, and SKF86002 were obtained from Sigma-Aldrich.

### Anti-p38α scFv selection by phage display

A large synthetic human scFv library developed in our laboratory was used to select specific anti-p38α scFvs by phage display according to a standard protocol^17,18^. Briefly, three subtractive rounds were first performed by incubating the library phages with p38β coated at 10 µg/mL on 96-well MaxiSorp immunoplates (Nunc) and by recovering the unbound scFv-phage clones at the end of each round. Subsequently, three rounds of selections were undertaken on p38α coated at 10 µg/mL.

### Production and purification of recombinant proteins

#### p38 production

All recombinant p38α proteins, wild-type or mutant, and p38β proteins were produced in BL21(DE3) as previously described^43^. Expression of GST- or 6xHis-fusion proteins was induced with 0.3 mM isopropyl β-D-1-thiogalactopyranoside (IPTG) at 30°C for 5h. Cells were harvested and frozen at -80°C. After thawing, pellets were resuspended in lysis buffer (10 mM HEPES pH8, 0.5 mM EDTA, 30 mM NaCl, 0.65% NP-40, 5 U/mL benzonase, 0.1 mg/mL lysozyme) and supernatants were clarified by centrifugation. For phosphorylated p38 (P-p38) production, p38 proteins were expressed in a BL21(DE3) strain transformed with a pACYC184 vector for the expression of untagged MKK6-EE. The obtained p38 proteins were highly phosphorylated and fully active^19^.

#### scFv production

ScFvs containing 6xHis and c-MYC or FLAG tags at their C-terminus were expressed in the periplasm of HB2151 cells after cloning into the pAB1 vector. Cells were grown in 2xTY medium with 100 µg/mL ampicillin (2xTYamp) at 37°C overnight. This culture was diluted 1:100 with 2xTYamp medium, and then grown at 37°C. When cultures reached OD_600_ = 0.5-0.7, 1 mM IPTG was added, and growth was continued at 30°C for 4 h.

#### Protein purification

After cell lysis, all proteins were purified by affinity chromatography on HisPur Cobalt resin (Thermo Scientific) or on Glutathione-Sepharose 4B (Pharmacia Biotech).

### Kinase assays

#### scFv inhibition of p38α phosphorylation by MKK6

Purified p38α (40 µg/mL) was activated by incubating with MKK6-EE (0.8 µg/mL) in the presence of scFv (80 µg/mL) and 200 µM ATP in kinase buffer (10 mM MgCl_2_, 0.62 mM EGTA, 0.25 mM DTT) at 22°C for 14h.

#### p38 kinase activity inhibition by scFv

The activity of P-p38 isoforms was tested using ATF2_(19-96)_ as substrate. ATF2 (2 µg) was combined with p38 (100 ng), scFv (1 µg), and 200 µM ATP in kinase buffer (final volume 15 µL). Reactions were incubated at 30°C for 30 min.

#### ADP-Glo assays

Except when specified for competition experiments, P-p38α (30 nM) or P-p38β (90 nM) were mixed with the peptide substrate IPTTPITTTYFFFKKK (SignalChem) (0.2 mg/mL) and different concentrations of the inhibitor, SB 202190 or scFv G9, in kinase buffer (40 mM Tris pH7.5, 20 mM MgCl_2_, 50 µM DTT, 0.1 mg/mL BSA). After 20 min pre-incubation at room temperature, ATP was added at the final concentration of 150 µM. Reactions were stopped after 60 min of incubation at room temperature, and ATP hydrolysis was measured using the ADP-Glo kit (Promega) following the manufacturer’s instructions.

The Bliss independent model was used to analyze the combination of scFv G9 and SB 202190. The two inhibitors were tested alone or mixed at different doses (detailed Figure S4A). The same concentration ratio corresponding to the ratio of their IC_50_ (150 nM for G9 and 30 nM for SB 202190) was maintained for all mixtures. The observed combination response was compared to the predicted combination response calculated with the data obtained with inhibitors tested alone at the same concentrations: (% p38α activity with [SB]) x (% p38α activity with [G9])/100.

### Epitope mapping

Deep Mutational Scanning (DMS) libraries were generated as outlined by Pruvost *et al*^24^. In summary, three distinct p38α DMS libraries were created: Library 1 encompassed amino acids M1 to Q120; Library 2 included K121 to G240; and Library 3 comprised T241 to S360. After library construction, yeast cells were transformed via gap repair, and transformation efficiency was assessed by plating serial dilutions on selective agar plates. Each library consisted of at least 5.10^5^ clones. Transformed cells were subsequently cultured and induced in galactose-containing medium prior to sorting.

After induction, yeast cells displaying the libraries were incubated in PBS-0.1% BSA solution containing either 20 nM G9 (biotin-labeled) and 28 nM B2-FLAG tag or 20 nM G9 (biotin-labeled) and 72 nM E5-FLAG tag. The incubation was performed under agitation for 2 hours at 20°C. Following this, cells were washed with ice-cold PBS-0.1% BSA, then incubated with APC-conjugated streptavidin and anti-FLAG tag/PE-conjugated antibody in PBS-0.1% BSA for 15 minutes on ice. Cells were subsequently washed with 1 mL of ice-cold PBS-0.1% BSA and sorted using a Cytoflex SRT cytometer (Beckman Coulter) with CytExpert software. Cells exhibiting decreased binding to one scFv while retaining binding to the other were collected. After sorting, cells were cultured at 30°C for two days in glucose-containing medium.

NGS sequencing and data analysis were conducted as previously described^23,24,44^. In summary, plasmid DNA from various yeast populations was extracted, followed by a two-step PCR to amplify the target region and incorporate Illumina adapters. Deep sequencing was performed using an Illumina MiSeq system (2x250 bp, v2 kit, 500 cycles), generating a minimum of 300,000 reads per population. Sequence reads were analyzed using custom scripts designed to trim, quality-filter, and count reads corresponding to each individual mutation in every sample. Enrichment scores were calculated by dividing the frequency of each mutant in the sorted population by its frequency in the unsorted population.

### Epitope validation

#### p38a mutation

Punctual mutations were introduced in p38α by site-directed mutagenesis of the p38α gene cloned in the pET vector, using the Kunkel method, as previously described^45^ (see specific primers in Supplementary Table S2). Recombinant proteins were produced and purified as described above.

#### Enzyme-linked immunosorbent assay (ELISA)

Nunc^TM^ Maxisorp 96-well plates (Thermo Fisher Scientific) were coated with purified wild-type or mutant p38α (1 µg/well). After saturation in PBS-3% milk at room temperature for 1h, c-Myc or FLAG tagged scFvs, diluted at the indicated concentrations in PBS/0.1% Tween-20/3% milk, were incubated at room temperature for 2h and then, detected with secondary HRP conjugated antibodies. After the last wash, HRP was revealed with 100 μL of TMB (1-Step Ultra TMB-ELISA, Thermo Fisher Scientific). Reaction was stopped with 50 μL of 1 M H_2_SO_4_ and absorbance was read at 450 nm.

### Cell culture

The THP-1, HeLa and HEK293 cell lines were cultured in Roswell Park Memorial Institute (RPMI) 1640 medium containing GlutaMAX (Gibco), 10% (v/v) heat-inactivated fetal calf serum and an antibiotic/antimycotic solution (Gibco).

### *In cellulo* antigen-antibody interaction assay

The human ORF sequences of p38α and p38β were cloned into the peGFP-C3 expression vector at the MGC facility (Biocampus, Montpellier) to encode p38α and p38β fused with the GFP fluorescent protein at the N-terminus. C-terminal NLS and NES tags were inserted in the resulting pCMV-GFP-p38 plasmids by circular polymerase extension cloning (CPEC). The pCMV-scFv-mCherry plasmids were generated by cloning the scFv coding regions into the pmCherry-N1 expression vector (Clontech) to encode scFvs in fusion with the mCherry fluorescent protein at the C-terminus.

The day before transfection, 10^5^ HeLa cells were seeded on coverslips. The plasmids (150 ng of pCMV-scFv-mCherry and 50 ng of pCMV-GFP-p38) were co-transfected using the Jet OPTIMUS transfection reagent (Polyplus), according to the manufacturer’s instructions. 48h after transfection, cells were fixed with 2% paraformaldehyde in PBS, permeabilized with 0.1% Triton X-100 in PBS, and DNA was stained using Hoechst 33358 dye (Sigma-Aldrich) at 5 µg/mL.

### THP-1 cell transduction for doxycycline-induced scFv-GFP-NLS fusion protein expression

The all-in-one autoinducible lentiviral vector pCLX-pTF-R1-DEST-R2-EBR65 (gift from Patrick Salmon; Addgene plasmid # 45952; http://n2t.net/addgene:45952; RRID:Addgene_45952) containing a TET-inducible promoter^46^ was used to clone B2 and G9 anti-p38α scFv genes in frame with the GFP gene and a NLS tag sequence.

Lentiviral particles were produced by co-transfection of the pCLX-scFv-GFP-NLS vector with the lentiviral packaging plasmid psPAX2 (gift from Didier Trono; Addgene plasmid # 12260; http://n2t.net/addgene:12260; RRID:Addgene_12260) and the VSV-G envelope expressing plasmid pMD2.G (gift from Didier Trono; Addgene plasmid # 12259; http://n2t.net/addgene:12259; RRID:Addgene_12259) into HEK293 cells as previously described^47^. Culture supernatants containing lentivirus particles were collected, filtered, and used for the infection of THP-1 cells. 48h post-infection, culture medium was replaced with fresh medium supplemented with 5 µg/mL blasticidin as selecting agent.

### THP-1 cell differentiation and LPS-induced TNF-α production

Stably transduced THP-1 cells were treated with 1 µg/mL doxycycline for 24h to induce scFv-GFP-NLS expression. As cells expressing G9-GFP-NLS, B2-GFP-NLS and GFP-NLS did not show comparable FACS signals, cells expressing the same level of GFP were selected with a FACS Aria® cell sorter (Becton Dickinson). Cells were immediately put back in culture and used for further experiments.

THP-1 transduced cells (10^5^ per well) were plated in 48-well culture plates and treated with PMA (10 nM) to induce differentiation into adherent macrophages^48^ and doxycycline (1 µg/mL) to induce scFv-GFP-NLS expression. After 48h, medium was removed, and cells were stimulated with LPS (5 ng/mL) for 4h. The secretion of TNF-α in culture supernatants was evaluated using ELISA MAX^TM^ Deluxe Sets (Biolegend) following the manufacturer’s instructions.

### Cytoplasmic and nuclear extracts

One million transduced THP-1 cells were treated with PMA and doxycycline for 48h as described above. After 30 min of stimulation with LPS (5 ng/mL), cells were washed twice with PBS, scraped and resuspended in 200 µL of hypotonic buffer (10 mM Hepes pH7.9, 10 mM KCl, 0.1 mM EDTA, 0.1 mM EGTA, and 1 mM DTT) and incubated on ice for 15 min. NP-40 was then added at a final concentration of 0.1%. Lysates were vortexed for 15 sec and centrifuged immediately (13,000 rpm, 15 sec) to yield supernatants containing the cytoplasmic fraction. Nuclear proteins were extracted by resuspending the pellets of nucleus in 30 µL of extraction buffer (20 mM Hepes pH7.9, 1 mM EDTA, 1 mM EGTA, 0.4 mM NaCl and 1 mM DTT). After 20 min incubation at 4°C under shaking, extracts were centrifuged (13,000 rpm, 5 min) and supernatants containing the nuclear fraction were collected. Both cytoplasmic and nuclear extracts were subjected to Western blot analysis with the indicated antibodies.

### Statistical analysis

Analysis of quantitative data was performed by using the Student’s t-test. Differences were considered statistically significant, when p < 0.05.

## Data availability

All the data are available in the main article and supporting information.

## Supporting information

This article contains supporting information.

## Supporting information

Supplemental Informations

## Acknowledgments

We thank the MGC, ARPEGE and PPM facilities (Biocampus, Montpellier) for plasmid constructions, time-resolved fluorescence analysis, mass spectrometry and SPR experiments respectively. We acknowledge the imaging facility MRI, member of the national infrastructure France-BioImaging (https://ror.org/01y7vt929) supported by the French National Research Agency (ANR-24-INBS-0005 FBI BIOGEN).

## Funding Sources

This work was supported by the Ligue Nationale contre le cancer comité de l’Hérault and by the French National Research Agency under the program “Investissements d’avenir” Grant Agreement LabEx MAbImprove: ANR-10-LABX-53. ER’s fellowship was funded by the Fondation ARC pour la Recherche sur le Cancer.

## Conflict of interest

The authors declare that they have no conflicts of interests.

## Abbreviations

The abbreviation used are: CD (common docking motif), CDR (complementary determining region), DMS (Deep Mutational Scanning), DSBU (disuccinimidyl dibutyric urea), FACS (fluorescence-activated cell sorting), GST (Glutathione-S-transferase), MAPK (mitogen-activated protein kinase), MKI (MAPK insert), MKK (MAPK Kinases), NES (nuclear export signal), NGS (Next Generation Sequencing), NLS (nuclear localization signal), PMA (phorbol 12-mystyrate 13-acetate), P-p38 (phosphorylated p38), scFv (single chain Fragment variable), SPR (surface plasmon resonance), LPS (lipopolysaccharide), TNF-α (tumor necrosis factor α), VH (heavy chain variable domain), VL (light chain variable domain), XL-MS (cross-linking mass spectrometry).

## Notes

### Competing Interest Statement

The authors have declared no competing interest.

