## Supplemental Informations for "Characterization of the p38α MAPK allosteric inhibition by a single chain Fv antibody"

[5] Deeptope SAS, Orsay, France.

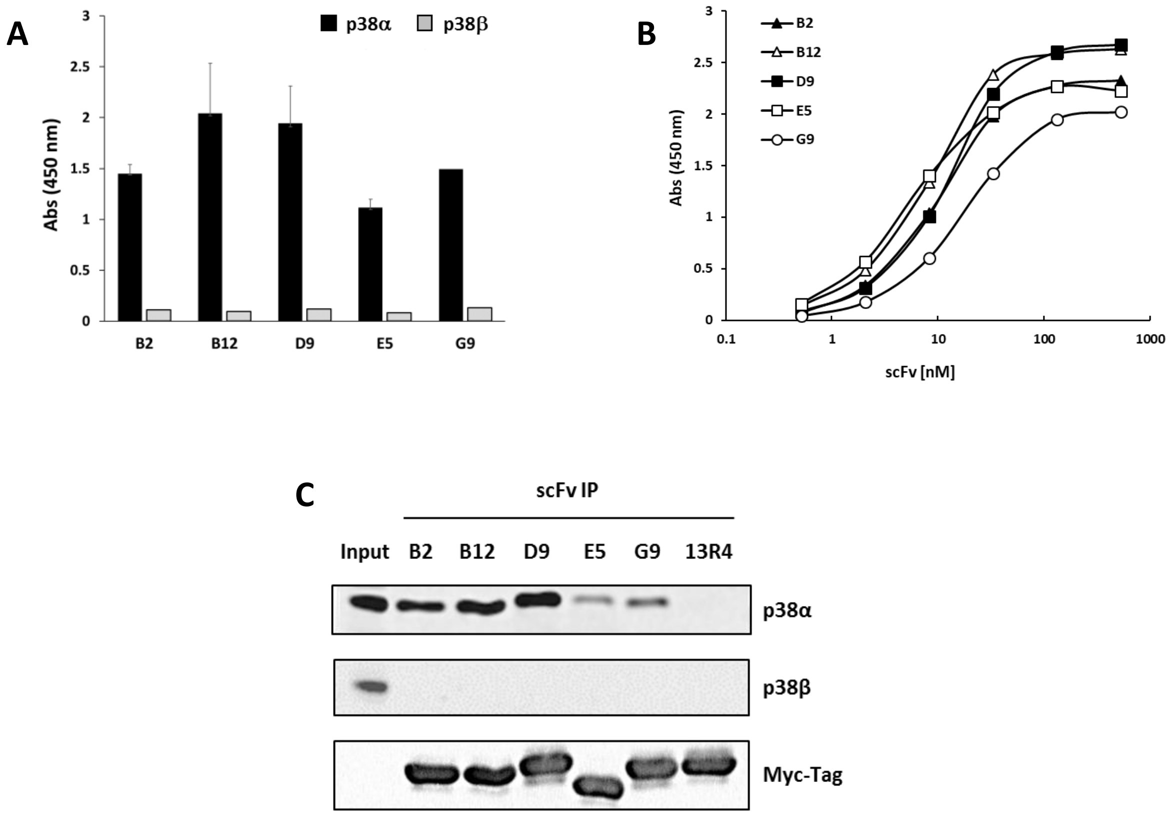


**Figure S1. Analysis of the binding specificity of five anti-p38α scFvs selected by phage display**.

**(A)** ELISA of unique scFv clones selected after three rounds of phage display panning. A total of 95 individual clones were selected from the polyclonal pool of panning round #3 for monoclonal ELISA assays against p38α and p38β. The positive clones were identified with a cut-off Abs(450nm) value >1, which corresponds to 5 times the background value. Sequence analysis of 19 clones with specificity for p38α revealed that five different scFvs were selected (B2 n=2, B12 n=3, D9 n=3, E5 n=2, G9 n=1). ELISA were performed with purified scFv produced in HB2151 periplasm. The binding of scFv was detected using an anti-MYC 9E10 antibody conjugated to HRP. Values represent means ± SD. **(B)** Dose-response binding of the selected scFvs to immobilized p38α. **(C)** Western blot analysis of p38α immunoprecipitation (IP). Purified scFvs, carrying 6xHis and MYC tags, were fixed on nickel beads and incubated with a total cell extract of HCT116 cells. The scFv 13R4 served as irrelevant antibody. Immunoprecipitates were probed on blots for p38α, p38β and MYC-tagged scFv detection.

**A**

| clones | H1 | H2 | H3 | L1 | L2 | L3 |
|---|---|---|---|---|---|---|
| **B2** | N**YN**MN | **S**I**R**GSSR**YIY**ADFVKG | S**SSNG**GMDV | AGTSSDVGG**GYN**VS | **N**DS**Y**RPS | SSYT**NY**STRV |
| **E5** | N**NS**MN | **S**I**R**GSSR**YIY**ADFVKG | S**SNDG**GMDV | AGTSSDVGG**GYS**VS | **N**DS**Y**RPS | SSYT**SY**STRV |
| **B12** | N**YY**MN | **G**I**Y**GSSR**GIN**ADFVKG | S**YY**GMDV | AGTSSDVGG**GSG**VS | **S**DS**S**RPS | SSYT**NY**STRV |
| **D9** | N**YY**MN | **G**I**Y**GSSR**YIS**ADFVKG | S**YY**GMDV | AGTSSDVGG**GSG**VS | **S**DS**Y**RPS | SSYT**NN**STRV |
| **G9** | N**YS**MN | **Y**I**G**GSSR**YIY**ADFVKG | S**SYYG**GMDV | AGTSSDVGG**YGG**VS | **Y**DS**Y**RPS | SSYT**YS**STRV |

**B**

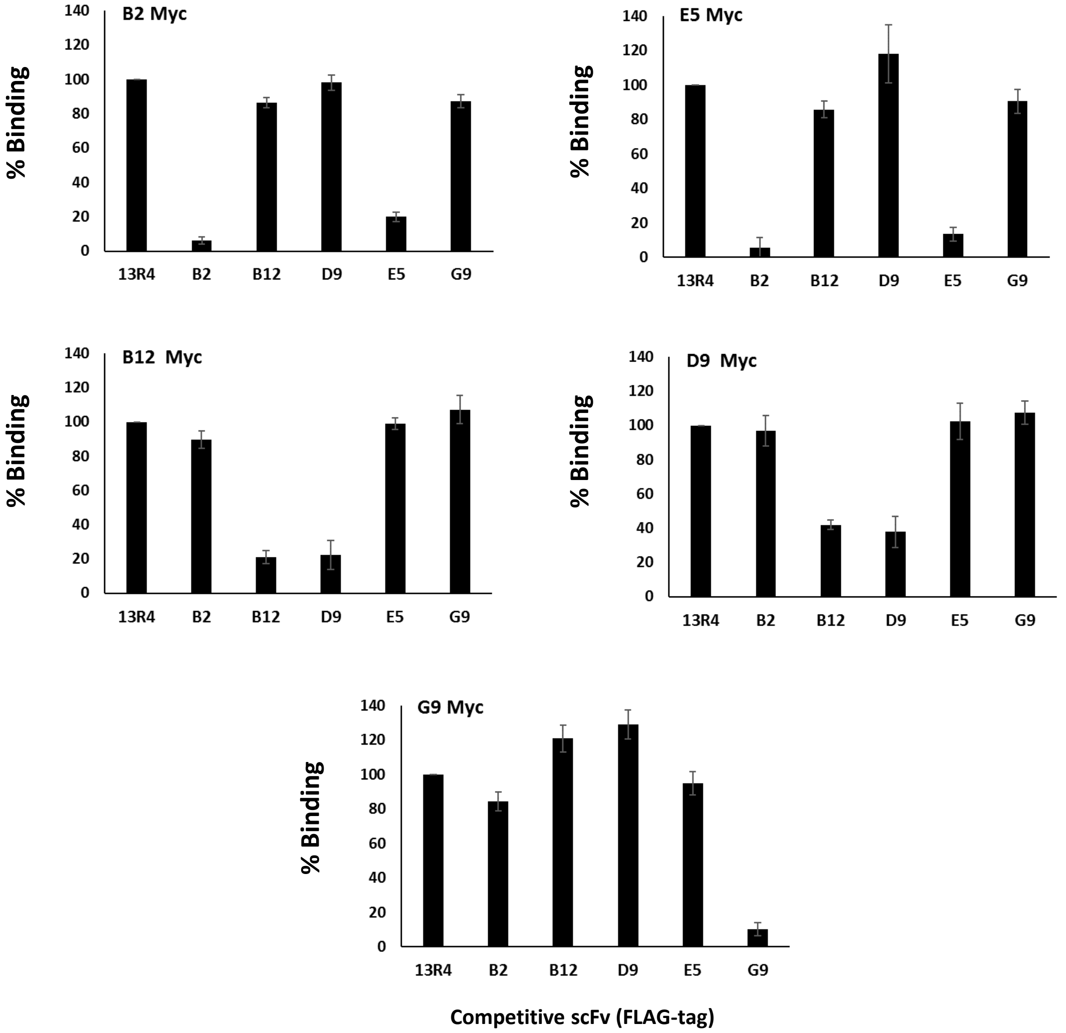


### Figure S2. Antibody-competition assay

**(A)** VH and VL CDR sequences of the anti-p38α scFv antibodies. Residue positions that are not modified in our scFv library, and are therefore found in every scFv, are indicated in grey. **(B)** Competitive binding assay. Competitive ELISA were performed by testing the binding of a MYC- tagged scFv (0.5 µg/mL) to immobilized p38α in the presence of competitor FLAG-tagged scFvs added at the saturating concentration of 50 µg/mL. The anti-MYC 9E10 antibody conjugated to HRP was used for detection. For each antibody, data were normalized to the control 13R4-FLAG competitor and presented as means ± SD from 3 independent experiments.

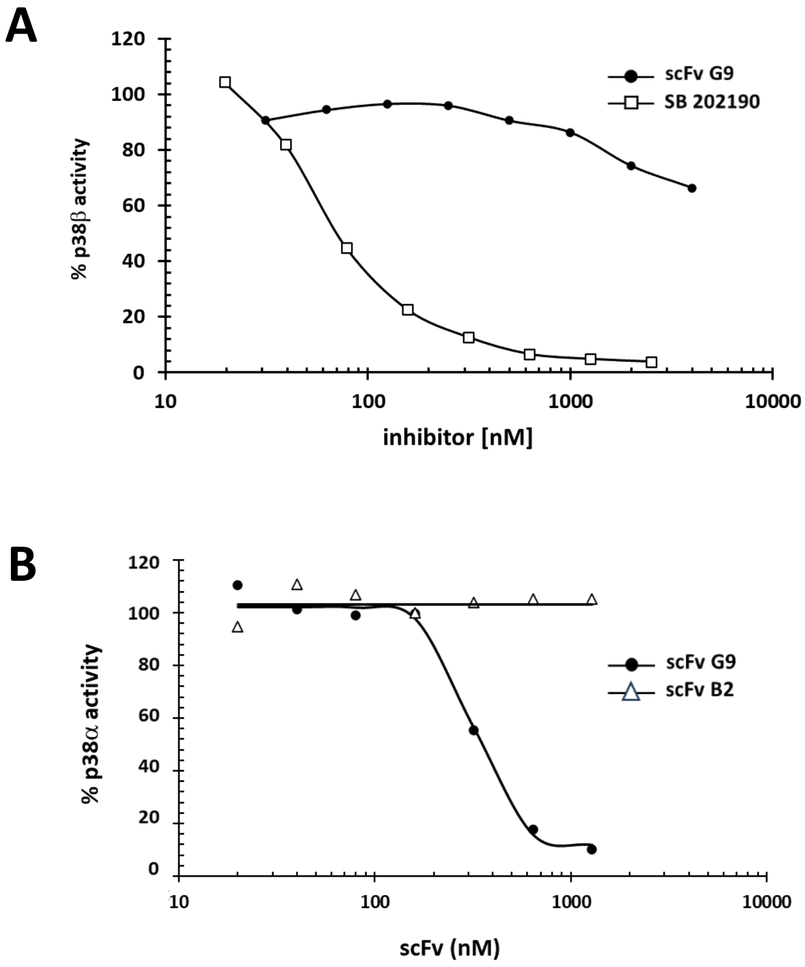


**Figure S3. Dose-response curve of p38α and p38β activity inhibition**

The kinase activity was analyzed using the ADP-Glo assay as described in Methods. (A) p38β inhibition by scFv G9 and SB 202190. (B) p38α inhibition by scFv G9 and scFv B2.

**A**

| | DOSE COMBINATIONS | | | | | | | |
|---|---|---|---|---|---|---|---|---|
| | 1 | 2 | 3 | 4 | 5 | 6 | 7 | 8 |
| scFv G9 (nM) | 0 | 18.75 | 37.5 | 75 | 150 | 300 | 600 | 1200 |
| SB 202190 (nM) | 0 | 4.375 | 8.75 | 17.5 | 35 | 70 | 140 | 280 |
| Ratio [G9]/[SB] | | 4.28 | 4.28 | 4.28 | 4.28 | 4.28 | 4.28 | 4.28 |

**B**

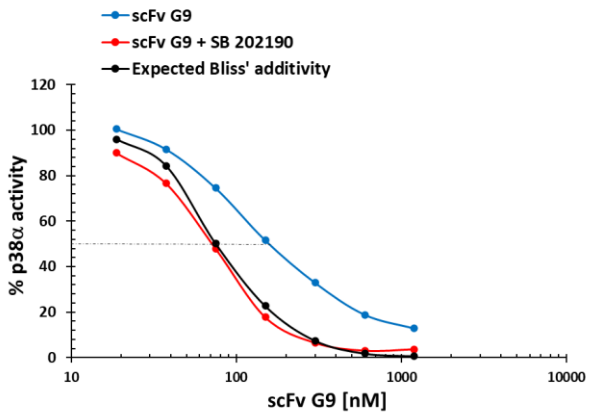


**C**

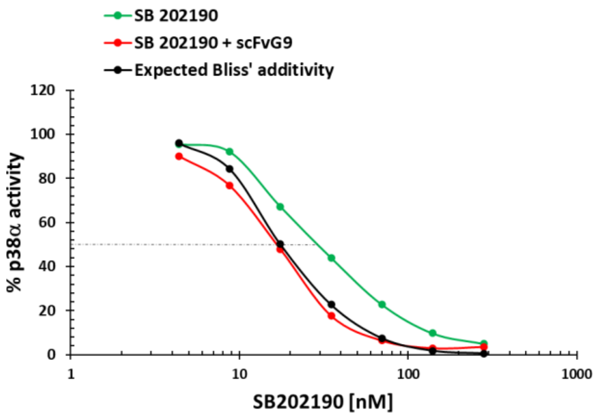


**Figure S4: Dose response inhibition of p38α activity by SB 202190 and scFv G9 tested alone or in combination.**

p38α kinase activity was measured in ADP-Glo assays using as substrate the peptide IPTTPITTTYFFFKKK. (A) Table indicating the different dose combinations of SB 202190 and scFv G9 tested. The mixture of the inhibitors at their $IC_{50}$ concentration (150 nM for G9, 30 nM for SB 202190) determined a concentration ratio which was then applied for each dose combination. (B and C) The black curve represents the percentage of p38α kinase activity calculated according to the Bliss' model assuming the independence of the two inhibitors.

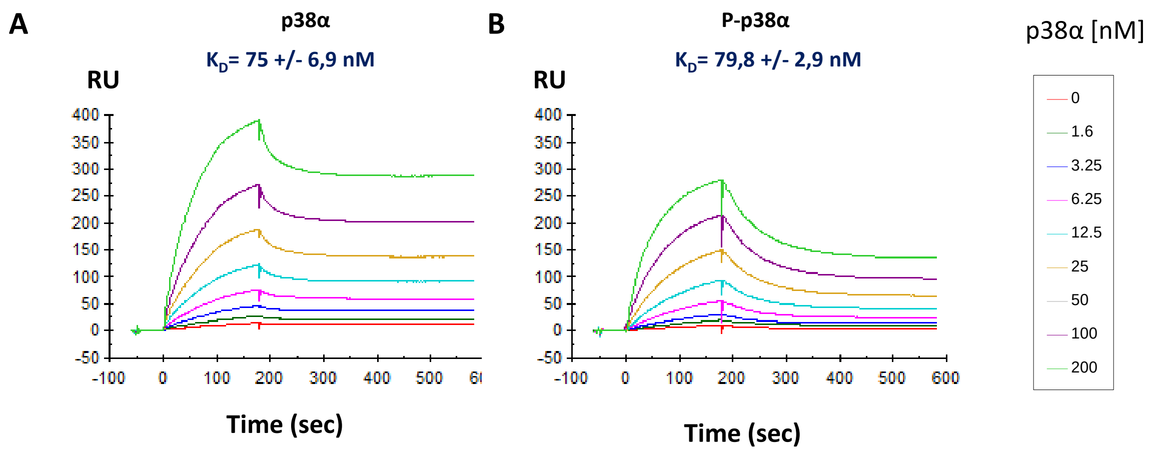


**Figure S5. Adjusted SPR sensorgrams analysing the binding of p38α or P-p38α to immobilized scFv G9.**

The antigen was injected at concentrations ranging from 0 to 200 nM.

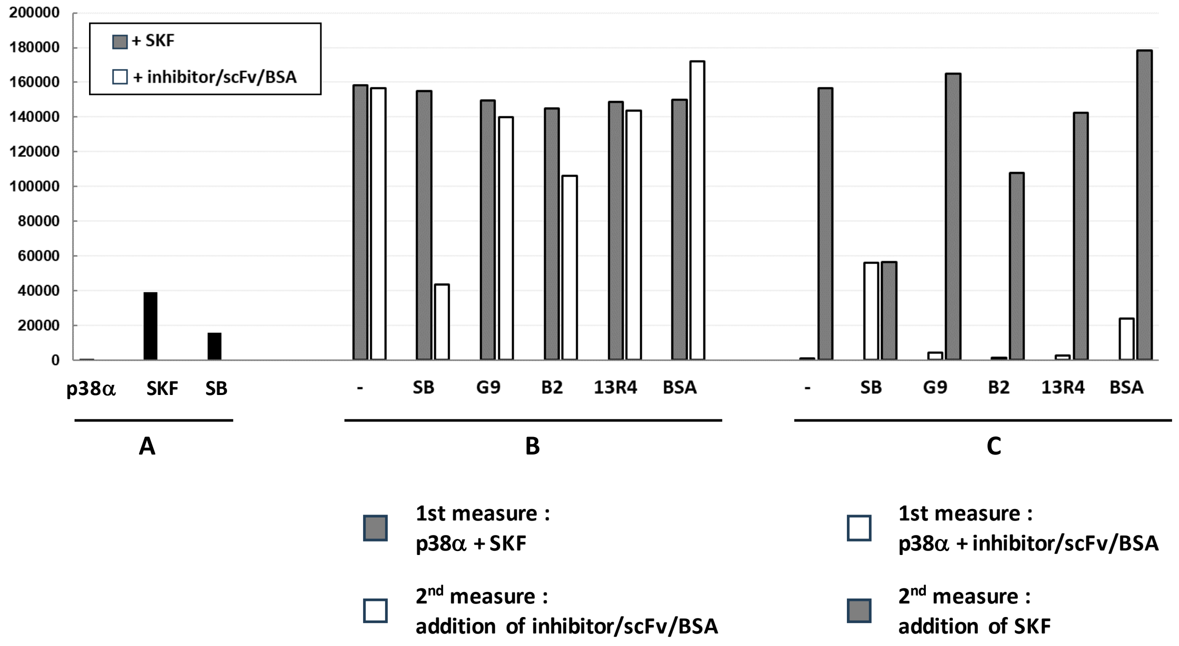


**Figure S6. Competitive-binding assay with the ATP-binding pocket inhibitor SKF-86002**

Endpoint analysis of SKF-86002 binding to p38α determined by measuring fluorescence emission at 420 nm (excitation wavelength: 340 nm). (A) Left panel: Histograms show individual fluorescence emission of p38α, SKF86002 and SB 202190 to indicate background of the experiment. (B) Middle panel: SKF-86002 was first added to a p38α solution and fluorescence immediately measured. Then, an inhibitor (SB202190, scFv G9) or a control protein (scFv B2, scFv 13R4, BSA) were added before fluorescence measured again. (C) Right panel: Fluorescence was measured before and after addition of SKF-86002 to a mixed solution of p38α with an inhibitor, scFv, or BSA. The concentrations used are: p38α 200 nM; SKF86002 200 nM; SB202190 2 µM; scFv 2 µM; BSA 2 µM. Fluorescence was monitored using the PHERAstar plate reader (BMG LABTECH).

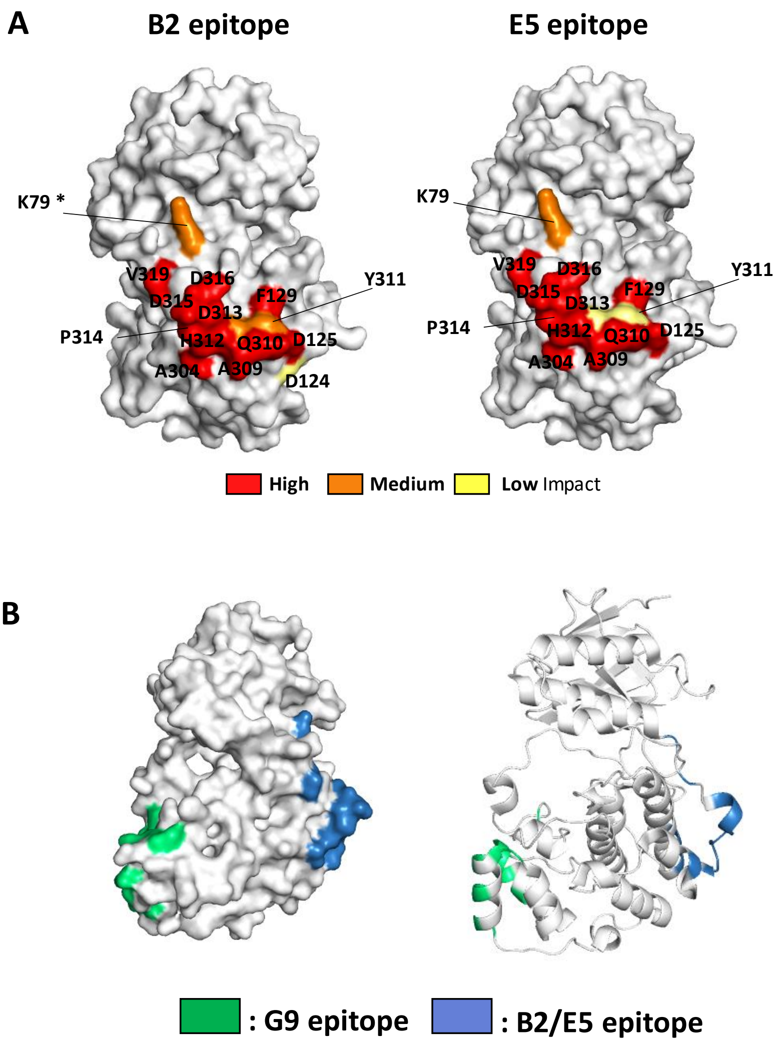


**Figure S7. scFvs B2 and E5 epitopes mapped by yeast display DMS approach.**

(A) Structures of p38α as a surface showing scFv B2 epitope (left) and scFv E5 epitope (right) (PDB 1WBT). The residues involved are colored from red (high) to yellow (low) to highlight their impact on the interactions. The asterisk (*) indicates K79 residue involved in DSBU cross-linking of scFv B2-p38α complex. (B) Structure of p38α as a surface (left) or a ribbon (right) showing G9 (green) and B2 (blue) epitopes, each located on an opposite side of the C-terminal lobe.

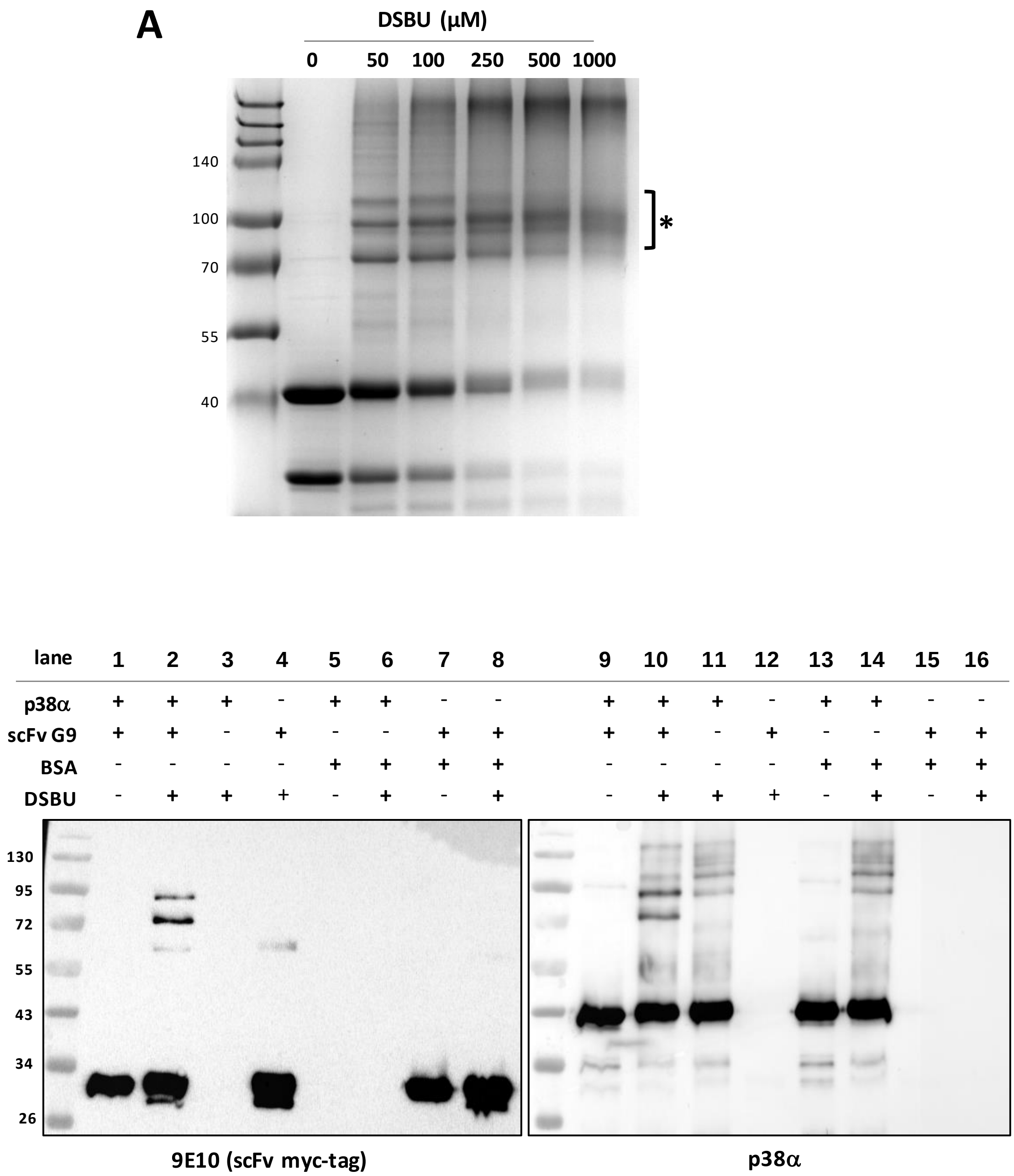


**Figure S8.** Identification of G9 binding site on p38α by cross-linking mass-spectrometry analyses.

**(A)** Optimisation of the DSBU concentration for p38α and scFv G9 cross-linking. A total of 10 μM protein was mixed at a molar ratio of 1:1 and incubated with various DSBU concentrations (0.05, 0.1, 0.25, 0.5, 1 mM) at room temperature for 1h. Cross-linking was analyzed by 10% SDS-PAGE and Coomassie blue-staining of the gel. *Band analyzed by MS-MS. **(B)** Western blot analysis of the cross-linking reactions using an anti-p38α antibody and the MYC-tag antibody 9E10 to detect G9. Reactions were performed as previously described using 5-fold molar excess of DSBU.

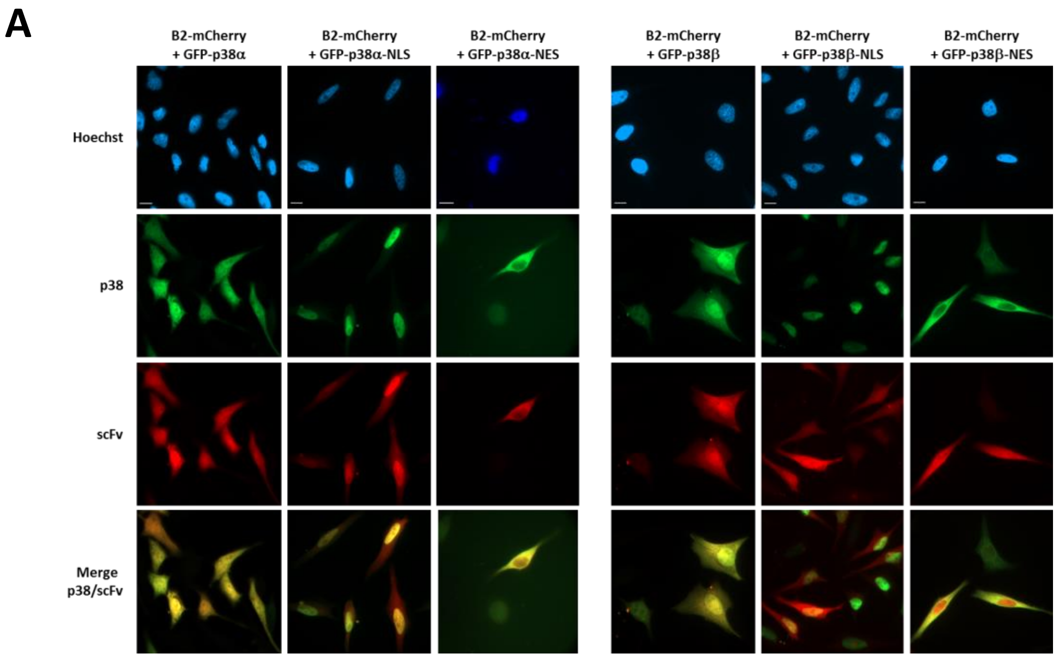


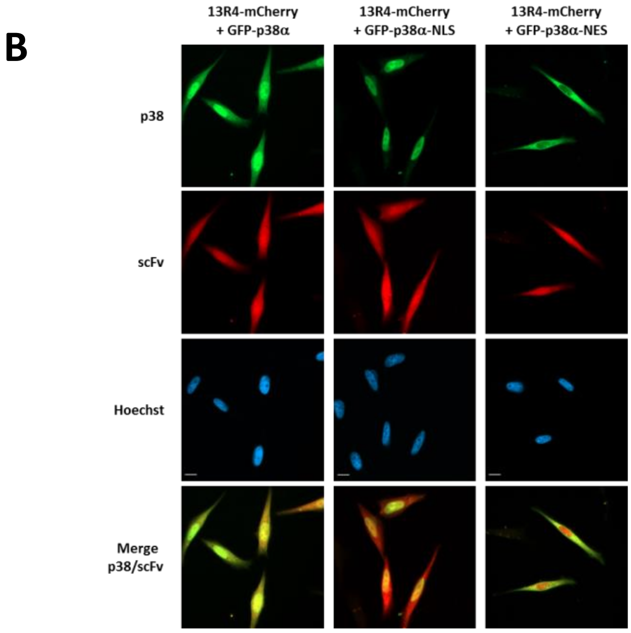


**Figure S9. Analysis of the binding properties of the anti-p38α B2 scFv in cells.**

Representative images of HeLa cells transiently transfected with the indicated scFv constructs tagged with mCherry and the indicated p38α constructs tagged with GFP. Cells were fixed 48h after transfection and the subcellular distribution of B2-mCherry (A) or 13R4-mCherry (B) proteins co-expressed with NLS- or NES-tagged GFP-p38α or GFP-p38β was visualized using a fluorescent microscope. The scFv 13R4 was used as negative control. Nuclei were stained with Hoechst. Magnification: x63.

Membrane 1

ponceau

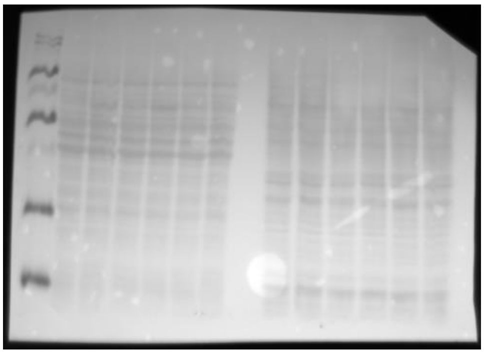

p38$\alpha$

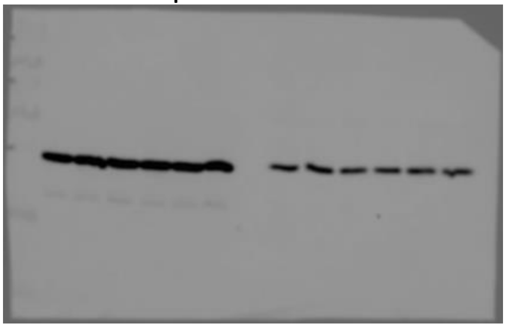

Phospho-p38

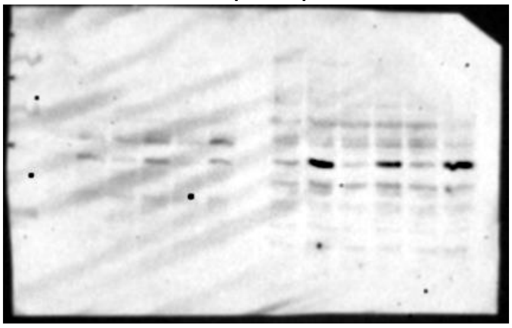

laminB1

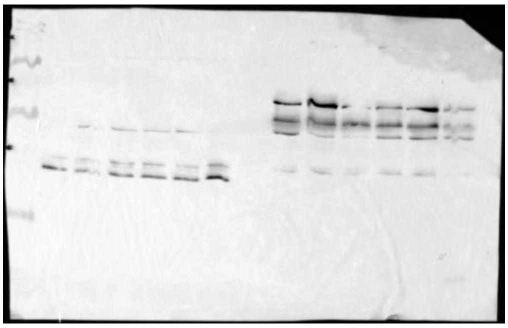

GFP

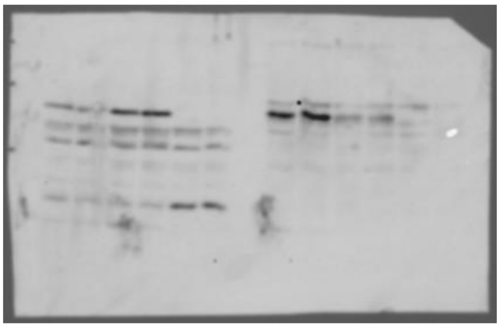

Membrane 2

ponceau

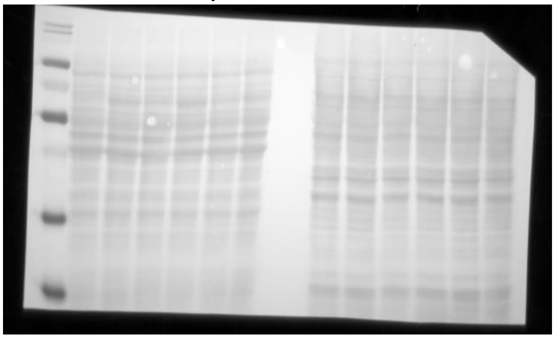

$\beta$-tubulin

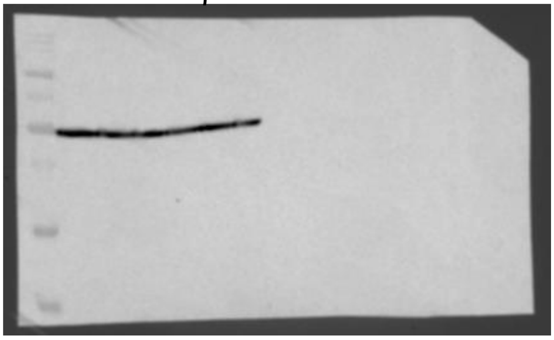

**Figure S10: Original Western blot images used for Figure 6B.**

**TABLE S1. Deep Mutational Scanning epitope maps of scFvs G9, B2 and E5**

For each scFv an enrichment score was calculated from the NGS results. The enrichment score represents the fold-change of the frequency of each mutation (left column) at every position of the antigen primary sequence (top row) after sorting compared to the unsorted p38$\alpha$ single mutants library. Enriched mutations are colored in red. On the bottom row the summary score is the number of mutations with an enrichment score > 1.5, for each position (index score >10 : key epitope residues; between 7 and 9 : medium importance; between 5 and 6 : low importance). No sort was performed with scFv G9 on the sub-library comprising residues 1 to 120 as no signal associated to a loss of binding could be detected. Gray columns represent a defect in the library, as no substitutions were generated at position P318.

**TABLE S2**

**Table S2. Primers used to introduce mutations**

| Mutation | Sequence 5' - 3' |
|---|---|
| T226S | GGTTCAGACCATATTGATCAGTT |
| H228Y | GGTACAGACTATATTGATCAGTTGA |
| Y258A | GCTATTCAGTCTTTGACTCAGAT |
| S261A | TATATTCAGGCTTTGACTCAGAT |
| D315A | AGATGAACCAGTGGCCGATC |
| V319A | TGATGAACCAGAGGCCGATC |
| AL for | GAGGAAATGACAGGCTACGT |
| AL rev | ATCTGCCTGCCGAGCCAG |

### Supplementary Methods

#### Surface plasmon resonance (SPR) binding affinity measurements

SPR experiments were performed on a T200 apparatus at 25 °C. The anti-c-MYC 9E10 antibody was immobilized on dextran CM5-S sensor chip by standard amine coupling according to the manufacturer's instructions. P-p38α and p38α proteins were injected at increasing concentrations (1.6-200 nM) on scFv G9 captured (around 400 RU) on immobilized 9E10 in the flow buffer (50 mM Tris-HCl, 100 mM KCl pH 7.5) containing 0.5 mM DTT. Gly-HCl (10 mM, pH1.7) was used as regeneration solution. All sensorgrams were corrected by subtracting the low signal from the control reference surface (without any immobilized protein) and buffer blank injections before fitting evaluation. The KD was calculated using a steady state fitting model (T200 Evaluation Software 3.0).

#### Cross-linking mass spectrometry

*Cross-linking reactions*. Reactions were performed using a freshly prepared stock solution of disuccinimidyl dibutyric acid (DSBU or BuUrBu; Thermo Fisher Scientific) in DMSO and added at a final concentration of 1 mM to a 10 µM protein solution containing 21 µg of scFv G9 and 29 µg of p38α diluted in 20 mM HEPES pH 7.6. Proteins were incubated at room temperature for 1h, and reactions stopped with Tris buffer (20 mM final concentration).

*In-gel trypsin digestion*. Protein extracts were separated by 10% SDS-PAGE and stained with Coomassie brilliant blue. Electrophoretic areas of interest containing G9-p38α complexes were cut and gel pieces (1 × 1 mm) were digested with trypsin according to a published protocol[38]. Briefly, gel pieces were washed in water, dehydrated with 50% acetonitrile (ACN) in 50 mM $NH_4HCO_3$, then 100% ACN and dried. After DTT reduction and iodoacetamide alkylation, gel pieces were re-swollen in 0.1 µg/µL trypsin (Promega) solution (100 mM $NH_4HCO_3$, 0.5 M $CaCl_2$, 1% proteaseMAX (Promega)). Resulting peptides were trapped and desalted on C18 Zip-Tips (Agilent) and speed-vacuum concentrated.

*Mass spectrometry analysis.* For LC-MS/MS, peptide mixtures were dissolved in 10 µl of 0.1% formic acid (FA) and loaded on an Ekspert 425 nanoLC system (Sciex) equipped with a C18 column (Discovery BIO Wide Pore, 3um, 0.5x10 cm, Supelco). The mobile phases were solvent A (water, 0.1% FA) and B (acetonitrile, 0.1% FA). Injection was performed with 98% solvent A at a flow rate of 5 µl/min. Peptides were separated at 30°C with the following gradient: 2% to 40% B in 105 min, 40% to 80% B in 5 min. The separation was monitored on-line on a TripleTOF 5600 mass spectrometer (Sciex). The total ion chromatogram (TIC) acquisition was made in information dependent acquisition (IDA) mode using

Analyst TF v.1.7 software (Sciex). Positive ions profiling was performed in the range from m/z 350 to 1500, followed by a MS/MS product ion scan from m/z 100 to 1500 with the abundance threshold set at more than 100 cps. The accumulation time for ions was set at 250 msec for MS scans and 100 msec for MS/MS scans. Target ions were excluded from the scan for 10s after being detected. The IDA advanced 'rolling collision energy (CE)' option was employed to automatically ramp up the CE value in the collision cell as the m/z value was increased. A maximum of 25 spectra per cycle were collected from candidate ions.

*Analysis of cross-linked peptides.* Peptides were identified using the ProteinPilot software v.5.0 (Sciex). For each MS2 spectrum, the Paragon algorithm[39] was used to search the Uniprot/Swissprot database with the following parameters: trypsin specificity, Cys-carbamidomethylation and search effort set to thorough parameters. Data were then converted into .mgf files to be processed by MeroX v.1.6.6 software to identify cross-linked peptides using the settings described elsewhere[27].
